# Cell-type specific impact of opioid use disorder and HIV on the human forebrain and cerebellum

**DOI:** 10.64898/2026.03.01.708876

**Authors:** Abbey A. Green, Tanmayi D. Vashist, Shweta Jakhmola, Xinze Chen, Gulshanbir Baidwan, Justin Buchanan, Shashi Kant Tiwari, Emily Griffin, Abigail Howell, Yuna Lee, David J. Moore, Sara Gianella, Davey M. Smith, Quan Zhu, Consuelo Walss-Bass, Allen Wang, Eran A. Mukamel, Kyle J. Gaulton, Tariq M. Rana

**Author notes:** These authors contributed equally Correspondence to T.M. Rana. These authors contributed equally (Abbey A. Green, Tanmayi D. Vashist, Shweta Jakhmola). These authors contributed equally (Xinze Chen, Gulshanbir Baidwan, Justin Buchanan).

## Abstract

Opioid use disorder (OUD), which frequently co-occurs with HIV infection, causes long-term neurological disease, yet the epigenetic and transcriptomic effects of OUD and HIV on specific cell types and regions of the brain are poorly understood. To assess the cell-type specific impacts of OUD and HIV across the human brain, we measured single cell transcriptomes and epigenomes of 580,353 cells in the prefrontal cortex, amygdala and cerebellum of 44 donors. We cataloged over 750k candidate *cis*-regulatory elements (cCREs) and identified gene regulatory networks (GRNs) of transcription factor activity across 35 neuronal and non-neuronal cell types. We identified specific neuronal and glial populations whose cCREs were significantly enriched for genetic risk of addiction-related traits. In OUD donors, we found evidence for reduced metabolic function in neurons in the PFC and cerebellum as well as increased gene expression related to voltage-gated calcium channel activity in the cerebellum. Using a cerebellar organoid model, fentanyl treatment reduced metabolic activity while increasing neuronal activity. Across brain regions, HIV activated immune-related pathways in glial populations, while comorbid OUD and HIV exacerbated metabolic changes in cortical glial cells. Cerebellum-specific Bergmann glia, in addition to forebrain microglia and astrocytes, showed expansion of reactive state identity in HIV. These results highlight shared and specific changes to immune, synaptic, and metabolic processes in OUD and HIV across brain regions and reveal that cerebellar cell types are distinctly affected by opioid abuse.

## Introduction

Opioid use disorder (OUD) is a major epidemic whose causes and neurodegenerative consequences are closely tied to reward and addiction-related circuits in the brain. While studies of addiction have focused on ventral midbrain regions^1,2^, addiction and neurodegeneration can impact forebrain circuits in the prefrontal cortex^3,4^ (PFC) and amygdala^2,5^ as well as the cerebellum^6–9^. OUD often co-occurs with chronic HIV infection, with long-term consequences that include cognitive decline and neurodegeneration^10^. Chronic OUD and HIV, acting over extended timescales of years to decades, may affect brain function in part by altering the epigenetic landscape of neuronal and glial cell types in the brain and consequently their transcriptional identity. Despite the important impacts of OUD and HIV on brain cells, cell type-specific data about the dysregulation of neuronal and glial identity in these disorders remains limited.

A fundamental challenge in understanding OUD and HIV neuropathology has been the cellular and molecular heterogeneity of the brain. Distinct neuronal and glial cell types express different receptor profiles, maintain different metabolic states, and have distinct circuit functions that confer differential vulnerability to drugs of abuse and viral infection. Opioids act primarily through the μ-opioid receptor MOR, encoded by *OPRM1*, which couples to inhibitory G proteins and broadly suppresses neuronal excitability and neurotransmitter release. Chronic opioid exposure induces neuroadaptations including receptor desensitization and downregulation, alterations in adenylyl cyclase signaling, and transcriptional reprogramming that collectively underlie tolerance and dependence. Many of these adaptations are encoded at the level of chromatin: sustained drug exposure reshapes the epigenetic landscape of neurons and glia, altering post-translational histone modifications and DNA methylation patterns that impact gene regulatory networks (GRNs) for months to years^11^. Similarly, infection by the retrovirus HIV establishes a latent reservoir in the central nervous system, integrating into the genome of microglia. Viral proteins including Tat and gp120 activate inflammatory signaling cascades that remodel chromatin accessibility across glial populations and drive persistent neuroinflammation even in virally suppressed individuals^12^. The cell type-specificity of these epigenomic changes, which differ between excitatory neurons, inhibitory interneurons, astrocytes, and microglia, cannot be resolved from bulk tissue analyses, motivating the need for single-cell approaches that jointly profile gene expression and chromatin accessibility across large numbers of donors.

The epidemics of OUD and HIV are closely linked. Opioid prescription for HIV-induced chronic pain elevates risk of OUD and, conversely, OUD-related needle use is a major risk factor for contracting HIV^13,14^. One third of individuals with OUD inject drugs, driving an increase in HIV infection^15^. Despite advancements in HIV treatment, modern long-term combination antiretroviral therapy (cART) does not fully protect the brain from the effects of HIV. HIV-associated neurocognitive disorder (HAND) occurs in 42.6% of the 37.9 million people living with HIV globally, including patients on cART.^16^ Transcription of HIV-encoded RNAs correlates with expression of genes related to inflammation, mitochondrial dysfunction, and neurodegeneration.^17^ Opioid use may amplify inflammation in those living with HIV^18,19^ and accelerate cognitive dysfunction^20^. Single-cell transcriptomic studies of OUD in the ventral midbrain and striatum have found elevated markers of DNA damage, neurodegeneration, and inflammation in glial cells^1,21^. The impact of HIV on microglia and astrocytes can be assessed by single cell transcriptomics^22,23^. In the ventral midbrain, a recent single-cell transcriptomic study investigated the interactions between HIV and substance use disorder^24^. Despite this, the transcriptomic and epigenetic changes across brain regions due to the interaction between OUD and HIV infection remain largely unknown. Specifically, although the cerebellum has been linked to cognitive processes related to addictive behavior^6^, the impact of OUD on cerebellar circuits and distinctive neuronal and glial cell types unique to the cerebellum, including Bergmann glial cells, has not been addressed.

The cerebellum is increasingly recognized as an active participant in reward, motivation, and addiction beyond its classical role in motor coordination^6,25,26^. It maintains reciprocal connections with key reward circuits (VTA, nucleus accumbens, PFC, amygdala) and sends direct glutamatergic projections to the SNc that elevate striatal dopamine levels^27^. Cerebellar Purkinje cells express functional D2 receptors that regulate social and reward behaviors independent of motor function^28^. The cerebellum and basal ganglia form an integrated network in which dysfunction at any node propagates system-wide^26^. Neuroimaging studies show cerebellar hyperactivity during drug cue exposure in opioid-dependent individuals^29^, and chronic opioid use is associated with cerebellar volume loss. The cerebellum contains unique cell types including unipolar brush cells and Bergmann glia^30^, absent from cortex and amygdala, suggesting that opioid- and HIV-induced changes there may be qualitatively distinct from forebrain effects. Despite this, whether OUD or HIV alters the transcriptional or epigenomic identity of cerebellar-specific cell populations remains entirely uninvestigated.

To examine the neuronal and glial impacts of OUD and HIV, alone and in combination, we performed single nucleus multiome (paired snRNAseq and snATACseq) assays in the PFC, amygdala, and cerebellum from 44 donors. We defined gene regulatory programs in neuronal and non-neuronal populations in each region and assessed changes in OUD and HIV, as well as OUD with HIV. We linked cell type-specific regulatory networks with genetic risk of addiction and other neurological traits. To validate our findings, we performed spatial transcriptomics on a subset of donors in the PFC and developed a cerebellar organoid model of OUD. Overall, our study uncovered widespread and region-specific changes in inflammatory, synaptic, and metabolic processes in OUD and HIV across diverse brain cell types. These findings provide insight into the molecular drivers of cognitive dysfunction in OUD and HIV and establish a foundation for developing therapies including those that span disease contexts.

## Results

### Cell type-specific gene regulatory maps of human prefrontal cortex, amygdala and cerebellum

To determine how OUD and HIV alter gene regulation across distinct brain regions and cell types, we profiled gene expression and chromatin accessibility from the PFC (n=24 donors), amygdala (n=16), and cerebellum (n=35) using single nucleus multiome (joint snRNA-seq and snATAC-seq) assays (**Fig. 1a**). In total, across all regions, we profiled 44 distinct donors including 9 non-disease, 7 OUD, 23 HIV+, and 5 OUD and HIV+ donors (**Supplementary Table 1, 2**). After removing low quality nuclei and doublets based on RNA and ATAC metrics (see **Methods, Extended Data Fig. 1c-f**), we clustered 123,270 high quality single nucleus profiles in the amygdala, 184,468 nuclei in the PFC, and 272,615 nuclei in the cerebellum (**Extended Data Fig. 1g**). We annotated the cell type identity of each cluster based on gene expression profiles of known markers^31–36^ (**Supplementary Table 3, Extended Data Fig. 1g**). We identified 35 major cell types across regions, including 12 inhibitory and 14 excitatory neuronal cell types as well as 9 non-neuronal cell types including microglia, astrocytes, oligodendrocytes, and oligodendrocyte precursor cells (OPCs) (**Extended Data Fig. 1h, i**). To make these data available for the research community we created an interactive data browser and analysis platform at https://brainome.ucsd.edu/SCORCH (**Extended Data Fig. 2**).

**Figure 1.**
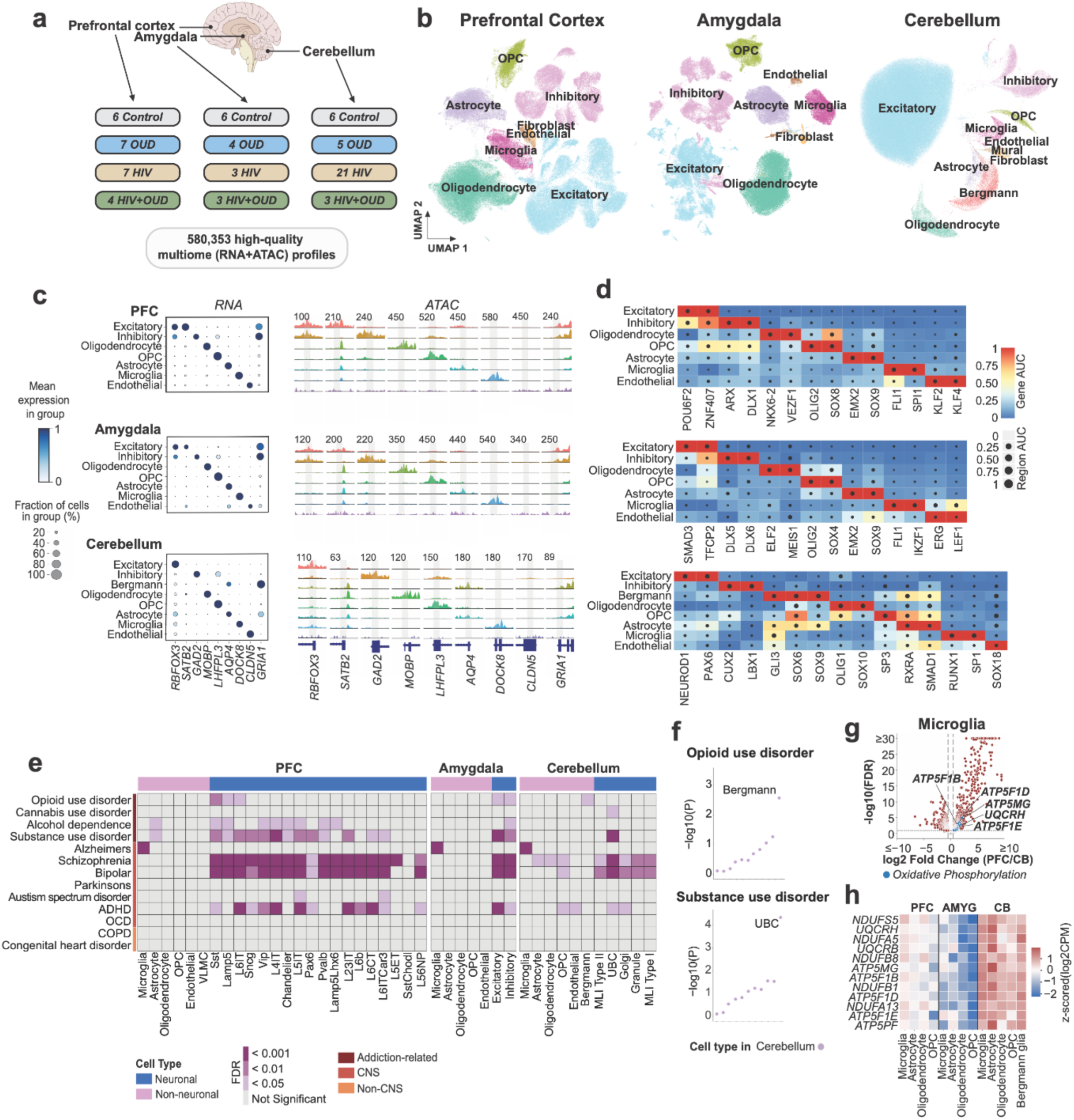
Overview of major cell types across brain regions and transcriptomic differences between regions in glial cell types. (**a**) Dataset overview. (**b**) RNA-based UMAP of all major cell types across brain regions. (**c**) Marker gene expression and genome coverage plots of major cell types across brain regions. Numbers represent track heights. (**d**) Up to two top GRNs ranked by cell type-specificity are shown for each broad cell type. Gene AUC represents enrichment of GRN gene targets in the most-expressed genes from each cell type. ATAC region AUC represents enrichment of GRN cCREs in each cell type’s most accessible ATAC peaks. (**e**) Linkage disequilibrium score regression (LDSC) showing significant enrichment of genetic risk in cell type-specific cCREs **(f)** Enrichment of genetic risk for substance use disorder and opioid use disorder in cerebellar cell type cCREs. Labeled cell types have significant enrichment of trait (FDR < 0.05). **(g)** Differentially expressed genes in PFC microglia compared to CB microglia. Red dots are significant (FDR<.10, |Log2FC|> .5) and blue dots are members of the oxidative phosphorylation pathway. **(h)** Normalized expression of oxidative phosphorylation pathway genes across glial cell types and brain regions.

To define the gene regulatory landscape of each cell type, we mapped candidate *cis*-regulatory elements (cCREs) across brain regions from control and disease donors and characterized their activity, cell type-specificity, and gene targets. We generated a consensus set of 757,121 cCREs across cell types from all three regions using MACS3^37^, which included both shared and cell type-specific peaks (**Extended Data Fig. 3**). There were on average 101,801 cCREs per cell type across all three regions. The majority of cCREs (96%) were non-promoter sites, and between 61-98% of the peaks in any cell type overlapped with previously reported bulk ENCODE cCREs^38^. Additionally, between 69-93% of the cCREs for any cell type overlapped with cCREs previously reported in the human brain *cis*-element atlas CATlas^39^. We used an activity-by-contact model^40^ to predict target genes linked to each cCRE for all cell types. The majority (60%) of predicted cCRE-gene links were cell type-specific, although related cell types such as OPC and oligodendrocytes shared more links (**Supplementary Fig. 2**). In addition, we defined genetic variants affecting cCRE accessibility by performing quantitative trait locus (QTL) mapping using RASQUAL^41^ (see **Methods, Extended Data Fig. 3**), focusing on inhibitory and excitatory neurons in each region (**Supplementary Table 5**). We identified TFs whose binding sites were enriched for chromatin accessibility QTL variants (**Supplementary Table 4**), including NFIX and ROR family TFs, which have roles in neuronal development and homeostasis^19,42–45^.

To identify transcription factors (TFs) and their associated target genes regulating cell type identity, we defined gene regulatory networks (GRNs) in each brain region using SCENIC+^46^. For each of 8 major cell classes, we identified the top two GRNs with the most cell type-specific target gene expression and chromatin accessibility (SCENIC+ RSS, **Supplementary Table 4, Fig. 1d**). These GRNs revealed key TFs involved in regulating cell type identity of these major cell classes. Several GRNs with cell type-specific activity were supported by previous literature identifying TFs driving cell type identity and fate, including PAX6 for excitatory cerebellar neurons^47^, OLIG1/2 for OPCs across all regions^48^, and DLX5/6 for inhibitory neurons in the amygdala^49^.

### Elevated metabolic gene expression in cerebellar glia

To understand variability in gene expression and regulation of similar cell types across brain regions, we examined glial populations in the PFC, amygdala, and cerebellum from non-diseased individuals. Glial cell types had highly conserved profiles across regions, where the correlation in gene expression profiles of the same glial cell type between regions was significantly higher (mean r=.93, SD=.007) than for different glial cell types within the same region (mean r=.85, SD=.007) (Mann Whitney U-test, p=5.14×10⁻^6^, two-sided) (**Extended Data Fig. 4b**). The majority of cCREs identified in each glial cell type in one region were shared with the same glial cell type in other regions (62-86%). We also found that similar TF binding motifs were enriched in cCREs of the same glial cell type across regions^50^ (FDR<0.1, **Extended Data Fig. 4c)**. For example, across all three regions, microglia cCREs were enriched for RUNX1 and ELF2^51–53^ motifs, astrocytes cCREs were enriched for RFX^54^ and SOX9^55^ motifs, and oligodendrocyte cCREs were enriched for SOX10^56^ and TCF12^57^ motifs.

A notable exception to the overall conservation of glial cell type-specific regulation across brain regions was a significant increase in metabolic gene expression in cerebellar glia compared to PFC and amygdala (**Fig. 1g-h**, **Supplementary table 5**). Genes with increased expression in cerebellar glia compared to other regions were enriched in metabolic functional pathways, including mitochondrial respiratory chain components such as *NDUFB1* and *NDUFS12* (**Supplementary Table 5, Extended Data Fig. 4e**). Cerebellum-specific Bergmann glia had higher levels of metabolic gene expression compared with glia in other brain regions (**Fig. 1h)**. Together, these results suggest that cerebellar glial cells may have greater baseline metabolic activity compared to PFC and amygdala glia, potentially reflecting differences in the relative proportion of glial and neuronal cells or in the level of neuronal activity between forebrain and cerebellum.

### Enrichment of neuropsychiatric risk variants at cell type- and brain region-specific regulatory elements

OUD has a strong genetic component, but the cell types mediating genetic risk of OUD and other addiction-related disorders across different brain regions remain undefined^58^. To link human forebrain and cerebellum cell types to risk of addiction or other neuropsychiatric disorders, we determined enrichment of disease-associated genetic variants from genome-wide association studies in cell type cCREs from each brain region using stratified linkage disequilibrium score regression (LDSC)^39,59^. Excitatory and inhibitory neuron cCREs in the PFC and amygdala were significantly enriched (FDR<0.05) in heritability for opioid, alcohol, and general substance use disorders (**Fig. 1d**). In particular, cCREs from GABAergic Sst and Lamp5 expressing interneurons, and from excitatory layer 6 intratelencephalic (L6 IT) neurons in the PFC were enriched for genetic variants associated with multiple addiction-related traits. In the cerebellum, unipolar brush cell (UBC) cCREs were significantly enriched for genetic risk for multiple addiction-related traits, and Bergmann glial cCREs were enriched for genetic risk of OUD (**Fig. 1f**).

Among neurological disease traits, we found that cCREs active in microglia across all regions were enriched for genetic variants associated with Alzheimer’s disease^60^. Regulatory elements in multiple neuronal cell types were enriched for schizophrenia and bipolar disorder-associated genetic variants. Notably, cCREs from cerebellar-specific UBCs were enriched for schizophrenia and bipolar disorder risk variants as well as for variants associated with addiction-related traits, suggesting a potentially broader role of these cells across neuropsychiatric disease (**Fig. 1f**). By comparison, variants linked with non-brain related traits such as chronic obstructive pulmonary disease (COPD) and congenital heart disease were not enriched. Overall, these results implicate specific cell types across brain regions in addiction and neurological and neuropsychiatric disease risk.

### Gene regulatory changes in neuronal cell types in the cerebellum and PFC in OUD

To understand how regulatory programs in the forebrain and cerebellum are altered in OUD, we identified differentially expressed genes in cell types from each region. There were considerable differences in the composition of neuronal cell types across donors from the amygdala, likely due to variability in the precise location of tissue dissections within the amygdala across donors. Thus, we focused neuronal OUD-related analyses on PFC and cerebellum. Deep layer excitatory neurons and Pvalb interneurons in the PFC, as well as cerebellar granule cells and interneurons, had the most differentially expressed genes in OUD compared to control (FDR<0.1, |Log2FC| > 0.5, **Fig. 2a, Supplementary Table 11**). Genes upregulated in neurons in OUD had highly region-specific effects, whereas downregulated genes were shared across neurons in both regions, including regulators of ATP metabolism such as *CKB* and *ATP5IF1* (**Fig. 2b, 2c**). Gene set enrichment analysis^61^ revealed that downregulated pathways (FDR<0.1) shared across all neuronal populations largely related to aerobic metabolism (**Fig. 2d)**, which included downregulated metabolic genes such as *COX5A*^62^ and *ATP5IF1*^63^ (**Fig. 2e**). Cerebellar neurons showed significant upregulation of genes related to voltage-gated calcium channel activity, GTPase-regulating processes, and cell-cell adhesion in OUD, which was not observed in PFC (**Fig. 2d,e**). Upregulation of voltage-gated/calcium ion channel-related processes was driven by increased expression of calcium signaling genes such as *CACNA1E, CACNB2, CACNA1B,* and *RYR1* across cerebellar cell types (**Fig. 2e**).

**Figure 2.**
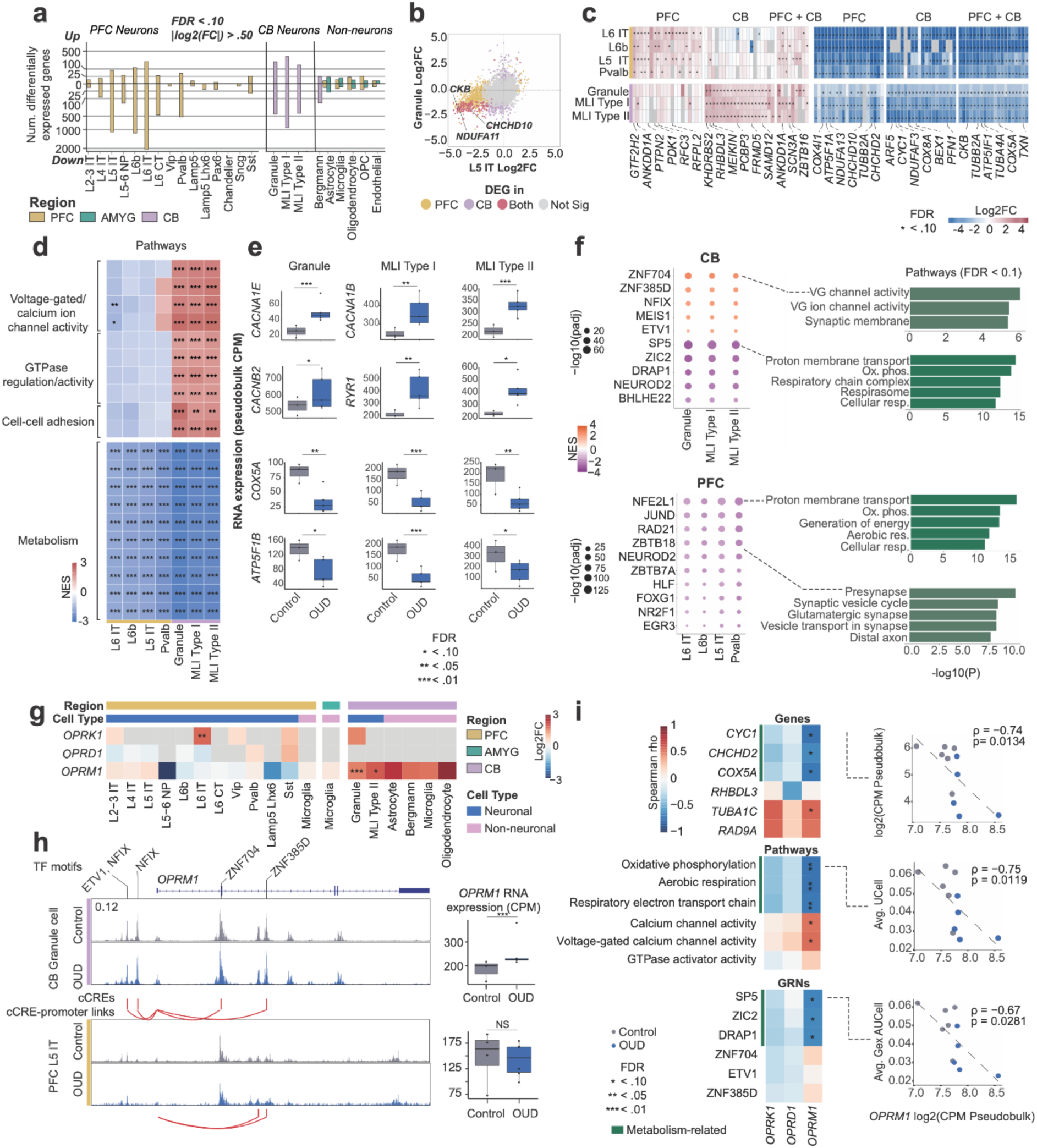
OUD-related impacts on brain cell types. **(a)** DEGs from OUD vs control analyses (FDR < .10, |Log2FC| > 0.5; PFC: n=7 control/4 OUD, amygdala n=3 control/4 OUD, cerebellum n=3 control/5 OUD). Females were excluded from CB DESeq2 analysis due to insufficient numbers. **(b)** Fold-change of gene expression in OUD vs. control donors in deep layer excitatory neurons of the PFC (L5 IT), and in CB granule cells. **(c)** Heatmap of expression fold-change for up to 20 up- (red) and down-regulated (blue) DEGs shared among PFC neurons (left), among CB neurons (middle), and across both PFC and CB neurons (right). Genes in grey were not tested for differential expression. (**d**) Top ten significantly upregulated pathways in OUD in CB neurons (top), and top ten downregulated pathways in PFC and CB neurons in OUD (bottom). **(e)** Expression of voltage-gated ion channel genes (*CACNA1E*, *CACNB2*, *CACNA1B*, *RYR1*) and metabolic pathway genes (*COX5A*, *ATP5F1B*) across conditions for cerebellar neurons. Boxes represent the interquartile range (IQR), whiskers represent 1.5 IQR, and the horizontal line indicates the median. **(f)** Gene regulatory networks (GRNs) enriched in OUD vs. control DEGs in neurons from the CB and PFC. Top 10 most significantly enriched GRNs shown for PFC. P-values for top significantly over-represented pathways are shown for select GRNs. (FDR < 0.1). VG = Voltage-gated **(g)** Heatmap of opioid receptor expression fold-change in OUD vs. control donors in cerebellar granule cells. **(h)** Open chromatin landscape showing ATAC-seq reads at the *OPRM1* gene locus in control and OUD cerebellar granule cells (top) and PFC L5 IT cells (bottom). cCRE are highlighted with pink vertical lines. TFs linked to peaks by DNA binding motif (SCENIC+) are indicated on top, and SCENIC+ links are indicated below. Right: expression of *OPRM1* gene. Boxes represent the interquartile range (IQR), whiskers represent 1.5 IQR, and the horizontal line indicates the median. **(i)** Heatmap of Spearman correlation between expression of opioid receptor genes in cerebellar granule cells, and cerebellar DEGs (top), metabolic pathways (middle), and metabolic GRNs (bottom) (n=6 control, 5 OUD donors).

To identify regulatory networks underlying neuronal responses to opioid use, we next assessed whether GRNs from our SCENIC+ analysis (described above) had altered activity in OUD. GRNs downregulated in cerebellar neurons in OUD (FDR<0.1) included SP5, ZIC2, DRAP1, NEUROD2, and BHLHE22, and the target genes of these GRNs were enriched for pathways related to aerobic metabolism. In PFC neurons, downregulated GRNs (FDR<0.1) included NFE2L1, RAD21, and NEUROD2 which were also enriched for aerobic metabolism-related pathways (**Fig. 2f, Extended Data Fig. 5g**). These results reveal specific TFs driving decreased expression of ATP metabolism processes observed in neurons across both cerebellum and PFC. In addition, several GRNs had upregulated activity in cerebellar neurons in OUD, including those regulated by ZNF704, ZNF385D, NFIX, MEIS1, and ETV1, and the target genes of these GRNs were enriched for voltage-gated ion channel activity and other synaptic transmission-related pathways (FDR<0.1, **Figure 2f, Extended Data Fig. 5g**). Interestingly, cerebellar neurons also had a smaller number of GRNs downregulated in OUD that regulated synapse-related pathways such as ZBTB18 and TFAP2A (**Extended Data Fig. 5g**), which may regulate multiple processes with both increased and decreased activity in OUD. In comparison, PFC neurons exclusively had downregulation of GRNs regulating synapse-related pathways, including those for ZBTB18 and JUND (**Figure 2f, Extended Data Fig. 5g**), supported by a decrease, although not significant, in these pathways in PFC neurons in OUD.

We next focused on the direct impact of opioid exposure on the brain by identifying cell types that express opioid receptors. Many excitatory and inhibitory neuron types in the PFC, amygdala, and cerebellum expressed the mu opioid receptor, *OPRM1*^58^, while a subset of neurons also expressed *OPRK1* and *OPRD1* (**Extended Data Fig. 4a**). Chronic opioid use may increase DNA methylation at the *OPRM1* promoter^64–66^, leading to downregulation of *OPRM1* expression in OUD. Interestingly, we found that *OPRM1* was significantly upregulated in cerebellar granule and MLI Type II cells in OUD compared to control donors (FDR<.10, |Log2FC|>0.5, **Figure 2g**). In contrast, *OPRM1* expression was not significantly altered in OUD across PFC cell types. Further supporting the increase in *OPRM1* in cerebellar neurons, cCREs at the *OPRM1* locus were more accessible in cerebellar granule cells than in PFC neurons (**Figure 2h**). Several of the upregulated GRNs regulating synaptic processes in cerebellar neurons targeted *OPRM1* (**Figure 2e,f, Extended Data Fig. 5g**). TFs regulating these GRNs were predicted to bind sequence motifs in cCREs linked to the regulation of *OPRM1* gene expression in the cerebellum using SCENIC+ (**Figure 2h**). These results highlight gene regulatory networks driving the upregulation of *OPRM1* expression and synaptic activity in cerebellar neurons.

We next investigated cellular processes associated with *OPRM1* expression in cerebellar neurons. We calculated the correlation between *OPRM1* expression and the expression of metabolism-related genes, pathways, and GRNs across control and OUD individuals (**Fig. 2i**). Overall, *OPRM1* showed significant (FDR<0.1), negative correlation with the expression of individual genes in metabolic pathways such as *CYC1, CHCHD2*, and *COX5A*. We used UCell^67^ to calculate gene signature scores for pathways altered in cerebellar neurons in OUD including metabolism and voltage-gated/ion channel pathways. Donor-averaged UCell scores for several metabolic pathways showed significant, negative correlations with *OPRM1* expression (Spearman r =−0.64, −0.67). Finally, donor-level gene expression AUCell scores for GRNs whose target genes were over-represented by metabolism-related pathways also showed significant, negative correlations with *OPRM1* expression (Spearman r=-0.69,-0.75). While other opioid receptors did not show a significant correlation with metabolism, these results highlight a consistent relationship between *OPRM1* expression and reduced expression of aerobic metabolism processes in granule cells (**Fig. 2i**). Of note, PFC neurons did not show significant upregulation of *OPRM1* in OUD, yet still exhibited down-regulation of metabolic processes, suggesting that these relationships may be more specific to the cerebellum.

Although OUD primarily impacted neurons, we also observed significant (FDR<.10) OUD-related changes in glial cell types across brain regions. Microglia have homeostatic and pro-inflammatory, disease-associated states. We annotated microglia states in each region based on reactive and homeostatic marker gene expression (e.g. homeostatic: *CX3CR1*, reactive: *SPP1*, see **Methods**) which revealed distinct populations (**Supplementary Table 21)**. Reactive microglia were significantly expanded in OUD donors compared to controls in the PFC and amygdala (Linear mixed-effects model (LMM), FDR<0.10, Coef = 3.295, 95% Confidence interval = .861-5.727, **Extended Data Fig. 5d)**, suggesting microglia contribute to neuroinflammation in OUD. Notably, we found many individual genes significantly downregulated in Bergmann glia in OUD (**Fig. 2a**), and both Bergmann glia and cerebellar oligodendrocytes showed significant downregulation of genes involved in metabolic pathways (**Extended Data Fig. 5f, Supplementary Table 12**). In addition, JAK-STAT signaling and T-cell activation-related pathways were significantly upregulated in PFC oligodendrocytes in OUD (**Extended Data Fig. 5f)**. These results suggest microglia, Bergmann glia, and oligodendrocytes may play a role in vulnerability to OUD in addition to neurons.

### Functional impact of opioid exposure in cerebellar organoids

Human donor studies are limited for studying complex disorders such as OUD due to heterogeneity across individuals including, for example, differences in drug dose and poly-drug use. We thus developed an *in vitro* cerebellar organoid (CERO) model from induced pluripotent stem cells (EC11) derived from primary human umbilical vein endothelial cells^68^, and utilized this model to validate the effects of opioids observed in the human cerebellum.

We observed CERO progressively across 3 months of *in vitro* maturation (**Extended Data Fig. 6a**). By day 16 *in vitro,* midbrain-hindbrain-specific markers appeared, including *EN2*, *PAX8*, *IRX3*, *GBX2* and *HOXA2,* while forebrain markers (*FOXG1*, *SIX3*, *OTX2*) were not detected (**Extended Data Fig. 6c**). At day 30 *in vitro*, we verified the presence of cerebellum-specific cells and their progenitors, including spatially organized *ATOH1+* rhombic lip (RL) and *KIRREL2+* ventricular zone (VZ) progenitors (**Fig. 3b**). Additionally, we observed *BARHL1+* granule cell progenitor cells (GCPs) and newborn *SKOR2+* Purkinje neurons (**Extended Data Fig. 6a**). After 3 months, we performed scRNA-seq and verified that CERO are composed of cerebellar cell types including granule cells, Purkinje cells, UBC, Bergmann glia, and other progenitor populations (**Fig. 3c,d**). The expression profiles of CERO cells were strongly correlated with the corresponding cerebellar cell types in primary tissue (granule cells: Spearman r=0.85, p<0.0001; Bergmann glia r=0.86, p<0.0001, **Fig. 3e**).

**Figure 3.**
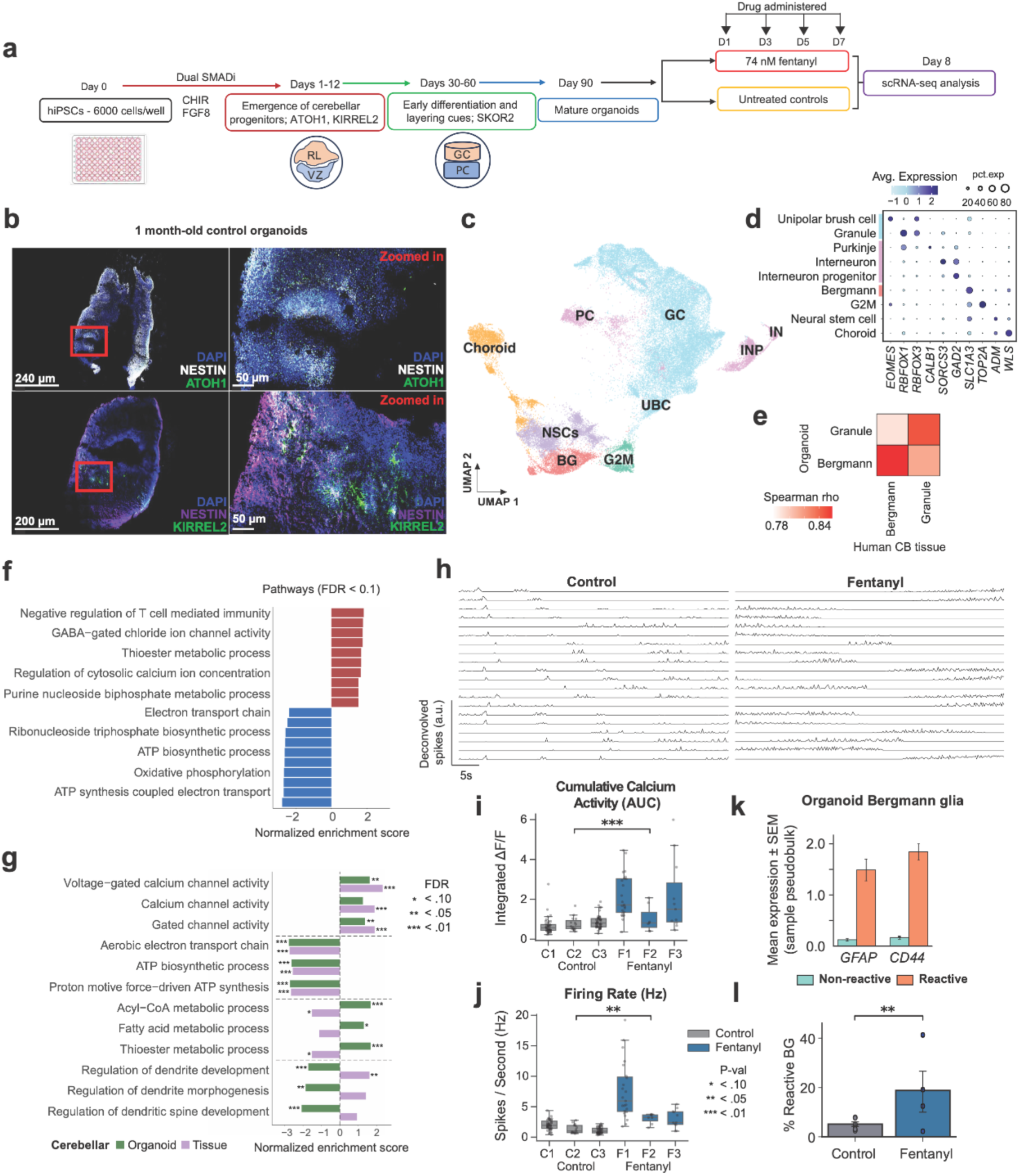
Altered metabolic pathways and activity in cerebellum organoid (CERO) model following sub-chronic fentanyl administration. (**a)** Experimental overview. (**b)** Immunofluorescence of ATOH1+ rhombic lip (RL) and KIRREL2+ ventricular zone (VZ) progenitors in 1 month old CERO. NESTIN identifies neural progenitors. **(c)** UMAP embedding of scRNA-seq profiles of nine major cell types identified in 90 DIV CERO. (**d)** Marker gene expression. **(e)** Heatmap of Spearman correlation between gene expression profiles of cell types shared in human tissue and CERO (**f)** Top 10 significantly up- and down-regulated pathways in CERO granule cells. Non-redundant pathways shown. **(g)** Shared and distinct pathway sets between human tissue and CERO granule cells. **(h)** Calcium-dependent fluorescence imaging from control (left) and fentanyl-treated (right) neurons. Each row shows the deconvolved spike rate estimated from one cell. **(i)** Cumulative calcium activity, calculated as the time-integrated normalized fluorescence (ΔF/F) for individual neurons in fentanyl-treated (N=3) and control (N=3) organoids. Data points represent single neurons (n=99 for control; n=41 for fentanyl-treated). Statistical significance was determined using a linear mixed-effects model (LMM) to account for organoid batch effects (***: p<.01). Boxes represent the interquartile range (IQR), whiskers represent 1.5 IQR, and the horizontal line indicates the median. **(j)** Estimated spike rate based on deconvolved calcium dependent fluorescence. Each data point represents a single neuron. (**: p<.05, LMM with organoid sample as a random effect). **(k)** Reactive marker gene expression in reactive vs non-reactive CERO Bergmann glia. **(l)** Proportion of reactive Bergmann glia, defined by expression of *GFAP* and *CD44*, in control and fentanyl-treated CERO. Error bars show standard error. **: p<.05.

To model changes in OUD, we cultured CERO with and without the opioid fentanyl every other day for seven days and performed scRNA-seq (see **Methods**, **Fig. 3a**). We then identified genes in each cell type with significant changes in expression in fentanyl-exposed CERO (N=4) compared to untreated CERO (N=6) (**Supplementary Table 13**). The effect of fentanyl on CERO gene expression was significantly correlated with the effect OUD on primary cerebellar granule cells (DEG Spearman r=0.23, p<0.0001), but not on Bergmann glia (Spearman r=0.04, p=0.59). We identified 113 pathways with significantly altered (FDR<0.1) expression in fentanyl-exposed CERO granule cells using gene set enrichment analysis (**Fig. 3f**). Granule cells in CERO showed similar changes in pathways as in granule cells from OUD donors, such as upregulation of calcium channel activity and downregulation of ATP metabolism processes (FDR<0.1, **Fig. 3g**). Despite the overall similarity of fentanyl-exposed CERO gene expression profiles to post-mortem human cerebellum, we also observed several differences (**Fig. 3g**). For example, processes linked to fatty acid metabolism and GABA/inhibitory synapse activity were upregulated by opioids in CERO granule cells but not in human cerebellum, while electron transport chain- and dendrite formation-related processes were suppressed in CERO granule cells (**Fig. 3f, g**).

To assess the functional consequences of fentanyl exposure in CERO, we performed live-cell imaging of calcium dependent fluorescence in fentanyl-treated (N=3) and control (N=3) CERO. These experiments confirmed the presence of functionally active neurons with distinct firing patterns across conditions (**Fig. 3h**, see **Methods**). Fentanyl-treated CERO showed significantly higher cumulative calcium activity (LMM, Coef = 1.069, p=0.001, 95% Confidence interval = .431-1.708, **Fig. 3i**) and firing rate (Hz) per neuron (LMM, Coef = 3.116, p=0.04, 95% Confidence interval = 0.127-6.105, **Fig. 3j**), which supports that fentanyl exposure increases transient calcium activity and enhances neuronal firing and excitability. Overall, these results validate transcriptome-level findings in CERO and primary granule cells by providing functional evidence that opioid use alters synaptic signaling in cerebellar neurons via increased transient calcium activity.

Bergmann glia react to injury and other insults^69,70^. Fentanyl is known to induce inflammation in microglia and astrocytes^71–73^, but it is unknown whether Bergmann glia similarly respond. In fentanyl-treated compared to control CEROs, we found the population of reactive Bergmann glia were significantly expanded (LMM, Coef = 1.13, p=0.019, 95% Confidence interval = 0.188-2.068) (**Fig. 3k,l**).

### Inflammatory gene expression in glia of HIV-infected donors across brain regions

Individuals with HIV have elevated markers of inflammation in microglia and astrocytes, even with viral suppression^74^. To examine how HIV alters gene expression in the cerebellum and forebrain, we performed differential expression analysis in cell types in each region in HIV donors compared to controls. Overall, we identified 1,568 unique genes with significant change in expression in HIV (FDR<.10) across regions, where the largest changes were in astrocytes and microglia. Microglial changes included downregulation of the homeostatic marker *CX3CR1* and upregulation of inflammatory cytokine *IL1B*. To validate our dataset against established findings, we compared our results to a previous study of PFC microglia; notably, genes upregulated in HIV donor PFC microglia in our cohort showed significant concordance, being significantly enriched for genes differentially expressed in HIV+ versus non-HIV+ populations in previous reports^74^ (*p* < .05, Fisher’s exact test, **Extended Data Fig. 7b**). A single HIV+ donor in our study was diagnosed with microglial nodule encephalitis, which can be caused by HIV, and had a strong signature of HIV-encephalitic gene expression in microglia (**Extended Data Fig. 7e**). Although we excluded this donor from analysis of changes in HIV, genes enriched in the encephalitic donor were strongly correlated with those upregulated in non-encephalitic HIV+ donors (**Extended Data Fig. 7f**).

Beyond microglia, HIV is known to alter gene expression in astrocytes^22^. However, whether the impact of HIV infection on gene expression is limited to microglia and astrocytes or extends to other glial or neuronal cell types across regions of the brain is unknown. The vast majority of changes in gene expression in HIV compared with non-disease donors were in glial cell types (91.7%, **Fig. 4a, Supplementary Table 11**). While changes in gene expression were driven primarily by astrocytes and microglia, other non-neuronal cell types including OPCs and oligodendrocytes contributed 22.9% of total DEGs. Changes in gene expression in glial cell types were strongly and significantly correlated across all regions (Spearman r=0.65-0.90, **Fig. 4c, Extended Data Fig. 7d**), and inflammatory pathways were consistently upregulated in glial cell types across brain regions in HIV+ donors. For example, the stress response gene *FKBP5*, which is upregulated with age in human brain cells^75^, was strongly upregulated (log2FC=2.08-2.5) in OPCs from HIV+ donors in all three regions. In contrast, synaptic pathways were selectively downregulated in glial cell types in the cerebellum. We found relatively few individual genes with significant changes in expression in neurons in HIV (**Fig. 4a**), although there were significant changes in metabolic pathways in neuronal cell types (**Fig. 4b**).

**Figure 4.**
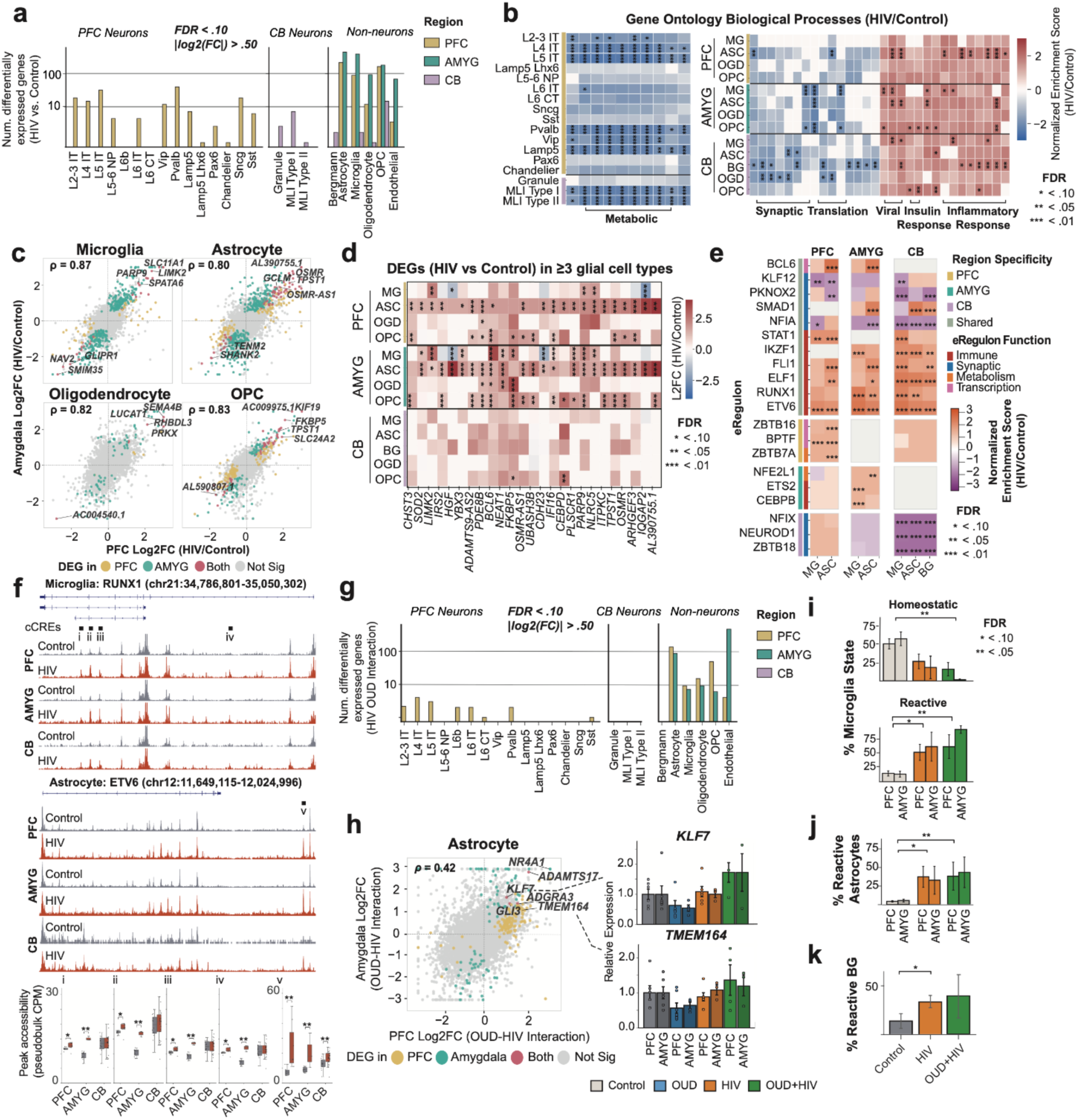
Overview of HIV and OUD-HIV interaction-related changes across brain cell types. (**a**) DEG count from HIV vs control analyses (FDR < .10, |Log2FC| > .50). Donor number for differential analyses: PFC (6 control, 6 HIV), amygdala (6 control, 3 HIV), cerebellum (5 control, 21 HIV). (**b**) Most common neuronal (left) and glial (right) gene ontology pathway terms from HIV/control DEG analyses. (**c**) Gene expression Log2FC correlation (HIV/control) between the PFC and amygdala across glial cell types. (**d**) Log2FC (HIV/control) of top DEGs occurring in 3+ glial cell types across brain regions. (**e**) fGSEA-based GRN enrichment (HIV/control). Normalized enrichment scores (NES) highlight GRNs significantly associated with HIV-responsive gene programs in glial cell types across brain regions. NES indicates up- or down-representation of GRN target genes. (**f**) Genome browser view (snATAC-seq pseudobulk) of HIV-responsive transcription factors RUNX1 and ETV6 in microglia and astrocytes. Gray shaded regions denote TF–binding motifs. Boxplots show pseudobulked region accessibility in highlighted TF-binding motifs. (**g**) DEG count (OUD:HIV interaction term, FDR < .10, |Log2FC| > .50). Boxes represent the interquartile range (IQR), whiskers represent 1.5 IQR, and the horizontal line indicates the median. Donor number for differential analyses: PFC (6 control, 7 OUD, 6 HIV, 4 OUD+HIV), amygdala (6 control, 4 OUD, 3 HIV, 4 OUD+HIV), cerebellum (5 control, 5 OUD, 21 HIV, 3 OUD+HIV). (**h**) Correlation in gene expression Log2FC (OUD:HIV interaction) between PFC and amygdala astrocytes (left). Expression of top correlated OUD:HIV interaction DEGs between PFC and amygdala astrocytes in donor pseudobulk, normalized to controls (right). Error bars show standard error. (**i**) Microglia state proportions across regions and conditions (Linear mixed-effects model, FDR correction). Error bars show standard error. (**j**) Reactive astrocyte proportions. (**k**) Reactive Bergmann glia proportions.

In addition to microglia, astrocytes have homeostatic and pro-inflammatory, disease-associated states^76–78^. We performed sub-annotation of astrocytes and microglia from each region based on reactive and homeostatic marker gene expression, which revealed distinct states of these cell types (**Fig. 4i, Supplementary Table 21)**. Reactive microglia and astrocytes were significantly enriched in HIV+ donors compared to controls in the PFC and amygdala (LMM, Microglia Coef = 2.145, Astrocyte Coef = 1.89, FDR<0.10, **Fig. 4i, j**). Likewise, reactive Bergmann glia in the cerebellum were expanded in HIV+ compared to control donors (LMM, Coef = 2.12, FDR=0.06) (**Fig. 4k**). Overall, these results support that reactive states of glial cell types across brain regions are expanded in HIV+ individuals.

To characterize the impact of HIV on GRN activity, we identified cell type-specific GRNs with target genes enriched for HIV-associated genes (**Fig. 4e)**. Target genes of RUNX1, associated with microglial activation^79^, and ETV6, which suppresses inflammation in a homeostatic feedback mechanism^80^, were significantly enriched in HIV-associated genes in glial cell types across brain regions (FDR<0.05) (**Fig. 4e, Extended Data Fig. 8b**). Supporting the increased activity of these GRNs in HIV, the expression of RUNX1 and ETV6 genes themselves were significantly higher in amygdala microglia (FDR<0.05, **Extended Data Fig. 8c**). Consistent with this, we found greater chromatin accessibility at enhancers targeted by ETV6 and RUNX1 in HIV donors (**Figure 4f**, **Extended Data Fig. 7d**). The target genes of RUNX1 and ETV6 included several immune mediators (*IL1B, JAK3, SPP1, PARP9*), and the effect of HIV on these genes was strongly correlated across brain regions (Spearman r=0.75-0.82; p<0.001, **Extended Data Fig. 8f-g**). Target genes of the ETV6 GRN were also enriched for HIV-associated changes in astrocyte and Bergmann glia (NES=2.1, FDR<0.001). Changes in ETV6 target genes in HIV were correlated across astrocytes in PFC, amygdala, and cerebellum (**Extended Data Fig. 8h-i**), as well as between Bergmann glia and amygdala astrocytes, suggesting ETV6 plays a cross-regional response to HIV in glial cell types. At the ETV6 locus, we identified a single cCRE linked to ETV6 that was significantly upregulated in astrocytes across all three brain regions (HIV/control: Log2FC=0.373-1.660, **Fig. 4f**), and this cCRE may help drive the increased activity of the ETV6-regulated network in HIV.

To determine whether HIV-associated gene expression changes were dependent on spatial context within regions of the brain, we performed spatial transcriptomics (ST) using multiplexed in situ hybridization to detect a panel of 266 genes including key neuronal and glial markers, as well as 9 HIV transcripts. After quality filters, we obtained spatial profiles in 102,253 cells in PFC of 2 HIV+ and 1 control donors, including one HIV+ donor with microglial nodule encephalitis. Detection of HIV transcripts in virally-suppressed donors is challenging due to low viral replication^74^ and limited sequencing coverage. We identified one HIV+ microglial cell located in the gray matter of the encephalitic donor, which expressed all 9 HIV transcripts (**Extended Data Fig. 7g**). We did not detect HIV transcripts in any other cells, indicating that HIV+ microglia are likely rare, or difficult to detect, even in donors with HIV-related encephalitic pathology^23^. We also found higher expression of reactive astrocyte marker *SERPINA3* in HIV donors (HIV+ Encephalitic: Log2FC=3.9, FDR <0.001; HIV+: Log2FC=2.5, FDR <0.001), particularly in white matter astrocytes (**Extended Data Fig. 7j**). By contrast, the antiviral response gene *IFITM3* was enriched in gray matter microglia (**Extended Data Fig. 7i)**.

### Astrocytes show cross-regional response to OUD-HIV interaction

The effects of OUD and HIV on the brain interact to exacerbate neurological disease^20^, although the genomic drivers of these interactions are poorly understood. We identified 194 genes where OUD significantly interacted with HIV to affect expression levels in PFC and amygdala astrocytes (OUD:HIV interaction term, FDR<.10, |Log2FC| >.5) (**Fig. 4g**), with fewer changes in other cell types. For example, genes including *KLF7* (FDR=.06,.07; Log2FC=1.02,1.7), *ADGRA3* (FDR=.03,.04; Log2FC=1.4,1.5), and *TMEM164* (FDR=.05,.11; Log2FC=1.15,1.55) were significantly upregulated by the interaction between OUD and HIV in both PFC and amygdala astrocytes (OUD:HIV interaction term, **Fig. 4h**). In addition to astrocytes, OUD and HIV interacted to affect endothelial cells. Notably, cytoskeletal gene *MAST4* was significantly upregulated by OUD-HIV interaction in both PFC and amygdala endothelial cells (Log2FC=2.33,3.28; FDR<0.10, **Supplementary Table 11**) yet was downregulated in OUD or HIV alone (|Log2FC|=1.17-2.14, FDR <.10).

We characterized biological processes altered in OUD and HIV interaction using gene set enrichment analyses. Metabolism- and translation-related pathways were significantly downregulated by OUD and HIV interaction across PFC glial cell types (FDR<.10, **Extended Data Fig. 9c**). We identified GRNs with target genes significantly altered by OUD and HIV interaction, and GRNs were most prominently affected in glial cell types. GRNs in glial cell types affected by OUD and HIV interaction in each brain region included NFE2L1 and RAD21, which were altered across glial cell types in both PFC and amygdala (**Extended Data Fig. 9c,e**). The target genes of NFE2L1 and RAD21 included *NDUFB9*, *NDUFA8*, and *NDUFC2,* which are all members of metabolic pathways broadly altered by OUD and HIV interaction (**Extended Data Fig. 9d**). These results suggest that OUD and HIV interact to disrupt the regulation of energy metabolism in cortical glial cells.

## Discussion

We generated an integrated single cell map of the human cerebellum, amygdala and prefrontal cortex across 44 individuals including those with OUD and HIV. These data provide an unprecedented resource of cell type regulation in specific regions of the brain, particularly in the amygdala and cerebellum which have been relatively unexplored by previous single cell studies^1,35,81,82^.

We found significant impacts of OUD and HIV in the cerebellum, including highly specific transcriptomic and epigenomic changes in cerebellar neurons in OUD relative to the other regions of the brain. Cerebellar neurons in OUD had elevated expression of voltage-gated and calcium channel pathways and gene networks, and cerebellar organoid models validated that opioid exposure increased both neuronal gene expression and activity. Prior evidence indicates that opioid use inhibits synaptic transmission by reducing voltage-gated calcium channel activity^83,84^, and thus the cerebellum may have unique neuronal responses to opioids. Increased activity of the opioid receptor *OPRM1* in cerebellar neurons correlated with pathways altered in OUD, and thus may play a role in OUD-related neuronal changes in the cerebellum. Genetic enrichment analyses further supported that cell types unique to the cerebellum such as Bergmann glia and UBC cells may mediate OUD effects. Overall, although a potential role of the cerebellum in OUD has not been widely investigated, these results highlight the cerebellum as a likely contributor to OUD pathogenesis and risk.

We propose that the cerebellum may contribute to OUD via at least two mechanistic routes. First, cerebrocerebellar loops connecting the cerebellum to the prefrontal cortex, striatum, and limbic system are known to modulate the timing, precision, and error-prediction signals that underlie reward learning^26,85^; chronic opioid-induced remodeling of activity in granule cells and Purkinje cells may therefore disrupt the reward circuitry’s ability to accurately predict and respond to drug-associated cues, reinforcing compulsive use^27–29,86^. Second, the cerebellum’s exceptionally dense granule cell population, which expresses high levels of *OPRM1*^6,7^, may be uniquely positioned to serve as a large reservoir of opioid-responsive neurons whose activity is sensitized by repeated drug exposure. The negative correlation between *OPRM1* expression and aerobic metabolic GRN activity in cerebellar granule cells suggests that sustained activation of mu-opioid receptor (MOR) couples to mitochondrial dysfunction in this population, potentially contributing to cerebellar atrophy as has previously been observed in an opioid rodent model^87^. Our data thus suggest a testable hypothesis for future mechanistic studies to examine whether cerebellar circuit manipulations can modulate opioid seeking behavior, withdrawal, or relapse.

Cell type-specific heritability analyses reveal that cerebellar Bergmann glia carry significant enrichments of genetic variants associated with OUD, positioning them as intrinsic biological substrates for susceptibility rather than passive downstream targets^27^. Direct cerebellar projections to the SNc modulate dopaminergic activity, and Purkinje cell D2R expression causally links cerebellar activity to reward-related behaviors^28^, suggesting genetic perturbations in these cell types alter reward circuitry through dopaminergic mechanisms. Upregulation of energy metabolism pathways in cerebellar glia compared to cortical and amygdala glia indicates these cells operate closer to their metabolic ceiling, rendering them vulnerable to energetic disruption from opioids or co-occurring viral pathology. Bergmann glia additionally display HIV-reactive transcriptional changes distinct from cortical astrocytes, extending cerebellar relevance to HAND. Together, these findings—genetic risk enrichment, elevated basal metabolic activity, and HIV-reactive Bergmann glia—support a model of compounded cerebellar vulnerability. Critically, upregulation of voltage-gated ion channel genes in cerebellar neurons under OUD conditions indicates neuronal hyperexcitability, a proposed biomarker for cognitive dysfunction^88^ and neurodegenerative progression^89^, raising the possibility that opioid-induced cerebellar hyperexcitability represents an early instigating event along a degenerative trajectory.

Beyond region-specific effects, OUD broadly downregulates aerobic metabolic processes across multiple brain regions, with opioid use shifting cellular metabolism toward glycolysis and disrupting mitochondrial function^90,91^ compromising energy-intensive processes such as synaptic transmission^92^. Reactive microglia were also expanded in OUD^73,93^ and opioid metabolites induce neuroinflammatory cytokines TNFα and IL-1β that elevate astrocytic glutamate release and synaptic activity^94^, together suggesting a state of bioenergetic exhaustion and dysregulation. This concurrent neuronal hyperexcitability and metabolic insufficiency aligns with models linking neuroinflammation-driven excitotoxicity to neurodegeneration^88,89,95^. The specific convergence of downregulated oxidative phosphorylation genes (*CYC1, CHCHD2, COX5A*) and upregulated voltage-gated ion channel activity in the cerebellum suggests this excitatory-metabolic imbalance is a region-specific feature of opioid neuropathology, potentially contributing to diffuse cognitive impairment in chronic opioid users and identifying mitochondrial support and excitatory tone modulation as therapeutic targets.

HIV broadly affects glial populations across brain regions, with increased proportions of reactive microglia, astrocytes, and Bergmann glia reflecting a state transition driven by viral neuroinflammation^78,96,97^. Shared upregulation of inflammatory pathways across brain regions and glial cell types, including oligodendrocytes and OPCs, highlights widespread glial vulnerability. Among the transcription factors mediating this response, ETV6—an ETS-family repressor that normally suppresses pro-inflammatory gene expression^80,98^—was upregulated in microglia, astrocytes, and Bergmann glia across all three brain regions, suggesting an attempted but insufficient compensatory response. Shared chromatin accessibility changes at ETV6 target enhancers indicate HIV establishes a convergent epigenomic inflammatory state across morphologically distinct glia. ETV6 target genes include key immune mediators *IL1B, JAK3,* and *SPP1*, and pharmacological strategies targeting ETV6 activity or its accessible enhancer locus in HIV+ astrocytes could represent a route to dampening the neuroinflammatory cascade driving HAND. The opposing regulatory relationship between RUNX1, which promotes microglial activation, and ETV6 suggests their balance may determine the magnitude and persistence of regional neuroinflammation in HIV.

A unique advantage of our study was the inclusion of donors with comorbid HIV and OUD, allowing us to investigate potential interactions between their effects. We found significant interactions, particularly in astrocytes in both the PFC and amygdala. The transcriptional interaction effects in astrocytes, including synergistic upregulation of genes such as *KLF7*, *ADGRA3*, and *TMEM164* in both PFC and amygdala astrocytes, suggest that opioids and HIV together engage a distinct astrocytic state that neither condition alone produces. Given that astrocytes coordinate synaptic glutamate clearance, blood-brain barrier integrity, metabolic support of neurons, and the propagation of inflammatory signals, a synergistic shift in astrocyte identity has the potential to amplify virtually every aspect of neurological impairment in comorbid individuals. The disruption of energy metabolism pathways in glial cells by the OUD-HIV interaction, driven in part through NFE2L1 and RAD21 GRN dysregulation and their mitochondrial complex I target genes, suggests a state of exacerbated metabolic failure superimposed on the individually deleterious effects of opioids and HIV. These interaction effects may help explain the accelerated cognitive decline and more severe neurological outcomes observed clinically in patients with comorbid OUD and HIV compared to either condition alone^20,99^, and identify astrocytic metabolic and transcriptional programs as priority candidates for therapeutic intervention in this high-risk population.

We note there are several limitations of our study for future studies to address. First, our cross-sectional postmortem design provides hypotheses for future causal studies, for example in cellular or non-human animal models. Inter-individual variability in drug exposure history, poly-drug use, comorbidities, postmortem interval, and cause of death may confound comparisons across donor groups despite our statistical covariate framework. Studies with larger numbers of donors will help to increase statistical power to detect subtle cell type-specific effects and improve the generalizability of interaction analyses. Our cerebellar organoid (CERO) model provides a controlled experimental system to validate fentanyl-induced transcriptomic and functional changes, but it cannot fully recapitulate the complexity of the adult human cerebellum, including mature circuit connectivity, glial-neuronal interactions, or the years-long exposure history characteristic of clinical OUD. Spatial transcriptomics in the PFC allowed us to identify spatial patterns of HIV-associated gene expression and localize the rare HIV+ microglial cell to cortical gray matter, yet the panel size and sequencing depth of the multiplexed in situ approach limited genome-wide resolution. Extending spatial analyses to the amygdala and cerebellum in larger cohorts will be important. Fourth, while we identified a high-confidence set of cCREs and GRNs, the functional validation of individual regulatory elements and transcription factor binding events particularly at the ETV6 locus and at OUD-associated GWAS-linked cCREs will require targeted experimental approaches including CRISPR-based perturbations in cellular models.

Overall, our cell type-resolved maps of the human cerebellum, amygdala and prefrontal cortex revealed widespread changes in OUD and HIV and highlight novel and unexpected roles of the cerebellum in these diseases. Our findings underscore the value of performing genomic studies across brain regions due to the highly region-specific effects of disease on cellular regulation in the brain. Moving forward, these data and results will help inform mechanistic studies aimed at understanding the cellular effects of OUD and HIV on the brain and the development of treatments for associated neurological impairments. Our findings converge on three major themes with broad significance. First, the cerebellum emerges as an underappreciated site of opioid-induced neuropathology, with cell type-specific changes in granule cells, Bergmann glia, and other cerebellar populations that are not observed or are of a qualitatively different character in the PFC and amygdala. This positions the cerebellum as both a contributor to OUD risk and a potential target for intervention. Second, HIV drives a convergent inflammatory state in glial populations across brain regions through shared transcriptional regulators, particularly the ETV6 and RUNX1 networks, providing molecular entry points for strategies to mitigate neuroinflammation in HAND. Third, comorbidity between OUD and HIV produces transcriptional interaction effects in astrocytes that go beyond additive effects, with particular impact on metabolic gene networks in PFC glial cells, underscoring the need to treat OUD-HIV as a distinct biological entity requiring specialized therapeutic strategies. Together, our dataset and visualization and analysis tools freely accessible at brainome.ucsd.edu/SCORCH provides a foundational resource to further dissect the molecular and cellular mechanisms of these intersecting epidemics and to identify novel targets for neuroprotective intervention.

## Methods

### Ethical statement

All research involving human postmortem brain tissue and human induced pluripotent stem cell lines was conducted in accordance with institutional guidelines and approved by the Institutional Review Board (IRB) at the University of California, San Diego.

### Post-mortem Multiome Analysis

#### Single nucleus multiome sample and library preparation

Snap-frozen aliquots of human brain tissue were provided for preparation of single-nucleus multiomic libraries using the Nuclei Isolation Kit (PN 1000447, 10X Genomics) with corresponding consumables (PN-1000448, 10x Genomics), Reducing Agent B (PN-1000450, 10X Genomics), and RNase Inhibitor Kit (PN-1000449, 10X Genomics) according to user specification. Briefly, nuclei were isolated from frozen tissue using the protocol for Single Cell Multiome ATAC + Gene Expression starting on page 33 of the user guide (<u>10X User Guide</u>, RevA, 10X Genomics) and nuclei were either manually counted on a brightfield microscope using a hemacytometer and Trypan Blue staining, or with an automated C100 Cell Counter (RWD Life Science) and DAPI staining.

Approximately 15,300 nuclei were used as input for profiling with the Chromium Next GEM Single Cell Multiome ATAC + Gene Expression kit (PN-1000280, 10x Genomics). All steps were performed according to user specification. Pre-amplification PCR was performed with 7 cycles, cDNA PCR was performed with 7 cycles. Indexing PCR for the ATAC-seq portion of libraries was performed with 7 cycles and sample indexes from the Single Index Kit N Set A (PN-3000427, 10X Genomics). 10 mL of cDNA library was used as input for library preparation of the RNA-seq portion; indexing PCR cycle number varied based on the input amount of library and per manufacturer specifications, falling in the range of 12-14 PCR cycles and using the Dual Index Kit TT Set A (PN-3000431, 10X Genomics).

QC sequencing was performed on the NextSeq 550 (Illumina) and Aviti24 (Element Biosciences) with a read scheme of R1:50, R2:50, I1:8, I2:16 (ATAC-seq) or R1:28, R2:91, I1:10, I2:10 (RNA-seq). Deeper sequencing to target 300M reads per library modality (300M for ATAC-seq and 300M for RNA-seq) was performed on the NovaSeq XPlus (Illumina).

#### Data processing

Raw RNA and ATAC sequencing data (FASTQ files) were processed using Cell Ranger ARC (v2.0.2, 10x Genomics) with a custom reference genome (GRCh38_HIV). Downstream analysis was performed using a custom multiomic pipeline, including preliminary quality filtering and removal of low-quality cells based on the number of RNA and ATAC fragments^100^. Following pipeline processing of each brain region, processed RNA data from individual samples were analyzed using Seurat5^100^ (v5.0.1). Data normalization and variance stabilization were performed using the SCTransform method implemented in Seurat, which accounts for sequencing depth and technical variation prior to downstream analyses.

#### Cell type annotations

Cell clusters were defined using the Leiden community detection algorithm^101^ based on principal component analysis-derived spaces from highly variable genes. Marker genes specific to known cell types were obtained from public databases and previous single-cell studies of similar cell types^31,35,81,102^. For PFC cell labeling, we used the Allen Brain Institute’s MapMyCells (RRID:SCR_024672).

#### Filtering by cell type

Distributions across percent mitochondrial reads, number of RNA fragments, number of ATAC fragments, ATAC mitochondrial reads, and TSS enrichment were determined for each annotated cell type. Barcodes within 2.5 median absolute deviations across these metrics were kept.

#### Transcriptome data analysis

Scrublet^103^ (v0.2.3) was used to remove doublets, a method that simulates doublets from the dataset and then computes the doublet score of each cell based on its similarity to simulated doublets. Poor quality cells were removed using control measures including gene count per cell and percentage of mitochondrial reads, which can indicate inefficient RNA capture or contamination respectively. To reduce data dimensionality and noise, principal component analysis (PCA) was performed by projecting genes onto the top 10-30 principal components. Cells were clustered using the Leiden community detection algorithm^101^ and results were visualized using uniform manifold approximation and projection (UMAP^104^ v0.5.5). From these clusters, we annotated cell types using published studies of human single cell transcriptomes from the respective brain region. Expression of the top 5-10 cell type-specific marker genes was used to annotate clusters of cells and validate predicted cell types using dot plots and violin plots to visualize marker gene distribution. We then created pseudobulk transcriptome profiles by summing the counts of all cells for each donor and each cell type. For example, Astrocytes_OUD1, Astrocytes_OUD2, Astrocytes_Control1. Pseudobulk profiles were used for analysis since it is the most rigorous and reproducible approach avoiding the potential for inflated significance when analyzing individual cell profiles^105^. Differential gene expression analysis was completed while accounting for potential confounding factors including sex and age using a negative binomial generalized linear modeling framework (DESeq2^105^ v1.42.0). Differentially expressed genes (DEGs) were defined using a false discovery rate (FDR) control procedure to account for multiple statistical comparisons. For HIV vs Control analyses, the design ∼sex + age + HIV was used, with only HIV and Control donors included. For OUD vs Control, the design ∼sex + age + OUD was used, with only males included and the sex covariate unused in CB and AMYG analyses due there being only one female donor. Only OUD and Control donors were included. To test the effect of OUD-HIV interaction we used the design ∼sex + age + HIV+ OUD + OUD:HIV and all donors were included.

#### Reactive glia analyses

To discern substates within glial cell types including microglia, astrocytes and Bergmann glia, we calculated the module score of substate marker genes. In microglia, P2RY12, CX3CR1, and TMEM119 were used as homeostatic marker genes, while HAMP, SPP1, TMEM163, and APOE^106^ were used as reactive markers. These marker genes then contributed to a module score used to separate homeostatic from disease-associated microglia. Several module-score thresholds were compared, and we selected the threshold (0.20) that provided the strongest separation of the reactive state based on marker expression, while retaining a sufficient number of reactive cells to avoid sparsity-driven noise. After removing donor pseudobulks with fewer than 50 microglia, less than 100 total microglia remained from both OUD and OUD+HIV donors in the cerebellum. Due to the low number of microglia in disease conditions, the cerebellum was excluded from this analysis.

For each donor, cell type proportions were calculated by aggregating cells across all samples and then transformed to the logit scale to generate a normal distribution. We modeled donor-level pseudobulk cell type proportions for each substate using a linear mixed-effects model. Fixed effects included condition, brain region, age (z-scored), and sex. Donor identity was included as a random intercept to account for repeated measures within donors. Significance of condition effects was tested by comparing each experimental group (HIV, OUD, OUD+HIV) to the control group within the mixed effects model framework. To control for multiple hypothesis testing across cell types, p-values were adjusted using the Benjamini–Hochberg false discovery rate (FDR) procedure, using a threshold of 10%.

Microglia state annotation methods were additionally applied to post-mortem brain tissue PFC and amygdala astrocytes, cerebellar Bergmann glia, and CERO Bergmann glia. Astrocyte reactivity marker genes included *CHI3L1, CD44, GFAP, SERPINA3, MAOB, MAPT,* and *SLC1A1*^77^, while reactive marker genes for BG include *GFAP*^69^ and *CD44*. In the CERO BG, we used *GFAP*^69^ and *CD44* as reactive markers. Because reactive marker gene expression was sparse in Bergmann glia in both post-mortem tissue and our organoid model, UCell^67^ was used to rank reactive marker gene expression rather than using the ScanPy score_gene function. For organoid Bergmann glia, a delta threshold of 0.05 was used for reactive annotation, while 0.25 was used for post-mortem tissue Bergmann glia and astrocytes.

For differential expression analyses, we created pseudobulk transcriptome profiles by summing the counts of all cells for each donor and each cell state. For example, Reactive_Astrocytes_OUD1, Non-reactive_Astrocytes_OUD1, Reactive_Astrocytes_Control1. Differentially gene expression analysis was completed while accounting for donor effects (∼donor + reactive_state) using a negative binomial generalized linear modeling framework^105^. DEGs were again defined using a false discovery rate (FDR) control procedure to account for multiple statistical comparisons. For microglia, the homeostatic population was used as a reference, while in astrocytes and Bergmann glia the non-reactive populations were used. Enrichr^107^ (gseapy wrapper, v1.1.2) was run using DEGs (FDR < 0.10, |Log2FC| > .5) and the GO Biological Processes 2025 library.

#### Permutation analysis of encephalitic HIV+ donor-associated genes in the PFC

Upregulated genes in the encephalitic HIV+ donor were determined by calculating the median difference in expression between all genes in this donor compared to controls, excluding one random control donor for validation. We took the top 200 genes with elevated expression in the encephalitic donor in both microglia and astrocytes separately, and computed the mean of the expression differences in these genes. This observed difference was then compared to 500 permutations of random 200 gene sets in a one-tailed permutation test across all conditions. The validation control donor was removed when creating the encephalitic donor gene set.

#### Gene regulatory networks (GRNs)

We used SCENIC+^46^ to infer enhancer-driven gene regulatory networks (GRNs), integrating single-nuclei RNA sequencing (snRNA-seq) and single-nuclei ATAC sequencing (snATAC-seq) data from our human postmortem brain samples. SCENIC+ was run separately for each brain region. GRNs were constructed per cell type, and condition-specific enrichment for GRNs was analyzed post hoc. Seurat’s subset() function was used to down sample the amygdala (123,270 subset to 76,282), prefrontal cortex (PFC 184,468 subset to 79,590), and cerebellum (272,615 subset to 74,992) multiome datasets, preserving even representations of all cell type and condition combinations while downsampling from the most abundant cell types first. SCENIC+ was run with the default parameters including nr_cells_per_metacells = 10, rho_threshold = 0.05, min_regions_per_gene = 0, and min_target_genes = 10.

#### GRN data preprocessing

We used preprocessed snRNA-Seq data from the upstream pipeline. Raw counts were stored in the adata.raw slot for downstream SCENIC+ analysis. For snATAC-seq data, we utilized pycisTopic^46^ to generate binarized topic models and identify reproducible accessibility patterns across cells. Topics above OTSU calculated thresholds were then retained. Quality control metrics included filtering cells based on total fragments, TSS enrichment scores, and fraction of reads in peaks (FRiP). Gene expression counts, chromatin accessibility, topics, and differentially accessible peaks across cell types were then used as input for SCENIC+ (v1.0a1).

#### GRN enrichment analysis (fgsea)

To identify pathways enriched in disease, we used the R package fGSEA^108^ (v1.30.0). Genes were ranked based on −log10(pval)*sign(log2FC) values from the DESeq2 differential expression results associated with the enrichment. Minimum and maximum gene set sizes were set to 15 and 500, respectively. Gene Ontology Biological Process (GOBP) gene sets were used for all pathway analyses. Molecular function gene sets (GOMF) were also tested for OUD^108^ control analyses to determine mechanistic functions related to channel and/or receptor activity.

To identify GRN target gene enrichment, we performed pre-ranked gene-enrichment analysis using the gsea R package^108^ with the fast multilevel permutation algorithm to test GRN target sets for coordinated regulation within each cell type–condition facet. GRN target gene sets were constructed from SCENIC+ direct GRN table: we read the output tab-separated file, split edges by GRN names so that each GRN corresponds to a unique set of target genes. For each cell type–condition facet, we restricted every GRN gene set to the overlap of genes present in the ranking vector for that pair and further enforced minimum/maximum gene-set sizes prior to testing. Per facet, genes were ranked from the DESeq2 results by signed –log10(p-val) score, 𝑠*_g_* = 𝑠𝑖𝑔𝑛(𝑙𝑜𝑔_10_(𝑝𝑣𝑎𝑙*_g_* + 𝜀)), with a small constant 𝜀 = 10^−300^. DESeq2 statistics were obtained from our pseudobulk differential expression result.

For each facet, we computed the running-sum enrichment score (ES) of each GRN using the fGSEA implementation on the pre-ranked list. We estimated nominal P values by permutations, then normalized the ES to obtain normalized enrichment score (NES) by dividing by the mean ES of size-matched random gene sets to correct for gene-set size bias. Multiple testing across GRNs within a facet was controlled by the Benjamini–Hochberg method. We recorded the leading-edge subset (the core group of genes contributing most to the ES) as defined in fGSEA, and we labeled the enrichment direction from the sign of NES (up = positive NES; down = negative NES).

#### Pathway over-enrichment in GRN target genes

We used fast over-enrichment analysis, which performs a hypergeometic test, to identify gene ontology pathways significantly over-represented within the target genes of a given SCENIC+ GRN. 10 and 500 genes were used as the minimum and maximum gene set sizes. All genes in SCENIC+ links for a region were used as the background/universe gene set.

#### Generating a cross-region consensus ATAC-seq peak set

tagAlign files were generated from CellRanger ATAC bam files for each single cell multiome experiment. A single, merged tagAlign file was then generated for each region. The tagAligns were then split by cell type using barcode-level assignments from snRNA clustering. Peaks were called for each cell type (including neuronal subtypes) using macs3^37^ with the parameters −q 0.05 —no model –keep-dup, using hg38 chromosome sizes for normalization. A consensus peak set was called across cell types from all brain regions, by identifying a representative peak with the highest pileup for a set of overlapping peaks. Peak widths were set to 501 bp by taking 250 base pairs flanking a given peak’s summit. Consensus peaks were mapped back to intersecting MACS3-called peaks to identify cell type peak sets. The average TSS enrichment (TSSe) across cell types was 3.15 (2.04-4.44), highlighting generally high quality epigenomic profiles.

#### Cell type transcription factor enrichment

We identified cell type-specific transcription factor enrichment for each cell type within each brain region to understand the regulatory programs driving cell type-specificity and function. We first calculated normalized (counts per million, CPM) cCRE-by-celltype pseudobulk profiles independently for each region. We then scaled the CPM values to represent probability values between 0 and 1 for each cCRE by dividing the counts in any given cell type by the total counts across all cell types. We then calculated the Shannon entropy for each cCRE using the Entropy() function from the DescTools package (v0.99.60). Low entropy peaks correspond to highly cell type-specific cCREs, i.e. cCREs whose amplitude is much larger in one or a few cell types compared to the rest.

For each cell type, we selected cCREs that had the maximum probability in that cell type. We selected the 2000 peaks with lowest entropy for each cell type. To identify transcription factor motifs enriched in cell type-specific cCREs, we used HOMER’s <u>findMotfisGenome.pl</u>, using all peaks detected across all cell types in this brain region as the background peak set, and a default region size as 200 base pairs.

#### Identifying differentially accessible regions

We implemented DESeq2^105^ to identify differentially accessible regions across different disease conditions. We prepared pseudobulk profiles of ATAC-seq peak counts for each cell type, which represent aggregated ATAC-seq peak counts across barcodes assigned to the cell type, for each donor. We tested peaks that have at least one count in at least 50% of the donors in each tested condition. DESeq2 was only run for cell types with greater than 500 barcodes. Age, sex, and mean cell type TSS enrichment were used as covariates, and continuous variables were scaled. As all OUD donors in our cerebellum dataset are male, we subset to using all males in our OUD vs. control DESeq2 analysis, removing sex as a covariate for these analyses.

#### Linkage disequilibrium score regression (LDSC) and cell type enrichment

We used LDSC to determine significant enrichment of heritability across multiple traits. We first selected the most well-powered, publicly available GWAS data sets across addiction-related traits, non-addiction central nervous system (CNS) traits, and non-CNS traits to serve as negative controls. We munged these GWAS statistics using munge_sumstats.py. Consensus peaks called in individual cell types were used to perform LDSC for each cell type, and LDSC was run separately for each brain region. Peaks were lifted over to hg19 and annotations of SNPs in provided cell type peaks were made using make_annot.py using the 1000 Genomes Project European population Phase 3 genotype data. We used the <u>ldsc.py</u> command to calculate LD scores, with 1000G_EUR_Phase3_basline and an LD window size of 1 cM. All regional peaks were used as an additional background set of peaks. Cell type enrichment was then calculated using the <u>ldsc.py</u> command with the –h2-cts flag, using the same baseline and background peak sets. BH multiple test corrections were applied across all the cell types tested within a region for each trait. Enrichments with FDR < 0.05 were considered significant.

#### Calculating correlations between opioid receptor expression and metabolism-related units in cerebellar granule cells

CPM-normalized pseudobulk counts for cerebellar granule cells were used to calculate the Spearman correlations between opioid receptors (*OPRM1*, *OPRK1*, and *OPRD1*) and other genes. Correlations were tested across the top 20 upregulated and top 20 downregulated DEGs shared by cerebellar neurons (ranked by number of cell types shared with, followed by minimum FDR across cell types). Several downregulated DEGs are components of aerobic metabolism pathways. UCell^109^ (v2.8.0) was run for the top 10 significant pathways up- and downregulated across cerebellar neurons (ranked by FDR), for granule cell barcodes. Pathway gene sets were filtered to only contain genes tested in DESeq2 in granule cells in OUD vs. control. UCell scores were averaged across barcodes from each donor and correlated with donor-level expression of opioid receptors. Finally, SCENIC+ gene expression AUCell values for cerebellar GRNs whose target gene sets are significantly over-enriched for metabolic and synaptic transmission-related pathways were donor-averaged across granule cell barcodes to correlate with opioid receptor expression. Units of correlation (DEGs, pathways, and GRNs) were not selected with a priori knowledge of correlation with *OPRM1*.

#### Genotyping

Genotypes for each sample were obtained through array-based sequencing with Illumina’s Global Screening Array (v3.0) and processed using GenomeStudio(2.0)^110^. Imputation was performed using the TOPMed R3 panel after which SNPs were filtered based on imputation accuracy scores (r2>0.7) using bcftools^111^ (v1.10.2).

#### caQTL Mapping

For single cell caQTL identification and analyses, we followed a previously described approach based on pseudobulk quantifications^112^. Cell type assignments for each barcode were used to split sample bam files into cell type-specific, sample-specific bam files using samtools^111^. The bam files were then used to make cell type pseudobulk count matrices using featurecounts^113^ (v2.0.0). A pseudobulk count matrix representing all peaks and barcodes was also generated to simulate bulk analysis. Count matrices were filtered for peaks containing an average of 5 reads per sample in the given cell type (or any cell type for bulk). Samples that were 3 or more standard deviations away from the mean in PCA analysis of the count matrices were removed from analysis for that cell type. PCA was run using all imputed variants that passed QC, and PCs 1-4 were used in covariate matrix to account for population structure. Covariate matrices for each cell type were generated using these genotype PCs, the principal components identified by the make_covariates function in rasqualTools^41^ (v0.0.0.9000), scaled age, and sex. Size factors were calculated and count matrices, covariate matrices and size factors were all converted to binary using rasqualTools.

Cell type and bulk tissue VCFs were made by merging VCFs from the appropriate samples and filtering for SNPs within 10kb of a peak in the count matrix. Allele specific counts were added to these VCFs from bam files using *createASVCF.sh* from RASQUAL. For each cell type, QTL mapping was performed with variants that intersected any of the cell type’s cCREs or a 10kb window around the cCRE.

#### Testing enrichment of caQTL variants that disrupt TF motifs

To examine transcription factor binding motifs disrupted by caQTL SNPs, we subset to lead SNPs for each caQTL. These SNPs were evaluated for disruption of human transcription factor motifs in JASPAR2022 from MotifDb (v1.44) using motifbreakR^114^ (v2.16) using the “ic” method. Enrichment of disrupted motifs in each cell type was measured using motifbreakR to identify JASPAR motifs from motifDB disrupted by any caQTL lead variant (lowest p value) within the caQTL feature regardless of significance. We performed a Fisher’s test on each motif in each cell type to identify motifs whose predicted disruption was enriched in variants with a significant caQTL association.

#### Identifying cCRE-gene links with Activity-by-Contact (ABC)

We used ABC^40^ to predict cCRE-gene links between peaks called and genes expressed in a cell type. tagAlign files split by cell type annotation were used as input. We used cell-type average Hi-C data provided by ABC with a Hi-C resolution of 5000. Peaks called by ABC were intersected with cell type peaks called using the previously used peak calling method used for snATAC data in this project. cCRE-gene links were additionally filtered to keep genes with pseudobulked CPM > 1 to account for genes expressed in each cell type. Self-promoters were removed from final cCRE-gene link sets.

### Organoid Data Processing and Analysis

#### Cerebellum organoid generation

The human induced pluripotent stem cells (iPSCs) EC11 derived from the primary HUVEC fetal cell line were differentiated to develop 3D cerebellar organoids^68^. The human-like cerebellum organoids (CEROs) showed laminar organization of various cell types specific to the cerebellum including Granule cell, unipolar brush cells and Purkinje cells. Briefly, iPSC EC11 were dissociated using accutase and resuspended in media containing growth factor reduced morphogen containing medium, composed of equal parts of IMDM (Gibco) and F-12 (Gibco), 5 mg/ml of high purity Bovine Serum Albumin (BSA) (Sigma), 15 µg/ml Apo-transferrin, 1% of chemically defined lipid concentrate (CDLC) (Gibco), and 450 M Mono-Thioglycerol (Sigma) with 7 µg/ml insulin (Sigma), 10 mM of SB431542, 50 ng/ml Noggin (Peprotech), 1.7 mM CHIR, and 20 mM Y-27632 ROCK inhibitor (Stem Cell Technologies). Approximately 6,000-10,000 iPSC EC11 in 100 µL per well of an ultra-low-attachment, V-bottomed, 96-well plates were added and allowed to aggregate. While maintaining a similar concentration of other morphogens the concentration of Noggin was increased to 100 ng/mL after 48hrs. To achieve that 50 µL of media containing 150 ng/ml Noggin was added after removing 40uL of the media in each well. On day 4, specification of midbrain hindbrain boundary was initiated by replacement of 40 µL media with 50 µL fresh media containing 200 ng/mL FGF8b and similar concentrations of SB431542, CHIR, and Noggin. ROCKi was no longer needed to achieve cell proliferation and aggregation and hence removed from the media. To further promote the development of cerebellar identity on day 6, 40 µL of media was replaced with 50 µL of fresh cerebellar differentiation medium (CerDM) I, comprising of DMEM/F-12 with 2 mM Glutamax (Gibco), 20% Knockout Serum Replacement (KSR) (Gibco), 7 mg/mL Insulin (Sigma), 15 mg/mL Apo-Transferrin (Sigma), 0.1 mM beta-mercaptoethanol (Gibco), 1X anti-anti solution (Gibco) supplemented with similar morphogens at concentrations used on day 4. On day 8, the final concentration of FGF8b was increased to 300 ng/mL by adding 100 µL of fresh media and removing 40 µL per well. On day 10, after removing 80 µL of media 100 µL of media maintaining a similar concentration of day 8 supplements was added. On days 12 and 14, a half media change was performed without changing the concentrations of any supplements by removing 70 µL of media and addition of 75 µL fresh CerDM1. The organoids were transferred to a 100 mm cell culture dish on day 16 and maintained under shaking condition (10 r.p.m) in CerDM II until day 30. CerDM II consist of DMEM/F12 medium (Corning), 1% B27 (Gibco), 1% N2 (Gibco), 2 mM Glutamax (Gibco), and 1X anti-anti solution (Gibco). A full media change was performed every 3 days. From day 30 onwards until day 60 the organoids were cultured in CerDM III, consisting of 1:1 DMEM/F12 (Corning) and Neurobasal medium (Gibco), 1%B27 with vitamin A, 1% N2, 2 mM Glutamax, 1% Chemically Defined Lipid Concentrate (Gibco), 5 mg/mL Heparin (Sigma) 1X anti-anti solution (Gibco), 0.5 ng/mL T3 (Sigma) and 1% Matrigel (Corning). The organoids were maintained in CerDM IV media (CerDM III lacking Matrigel and enriched with 14 ng/ml Brain Derived Neurotrophic Factor (BDNF) from day 60 onwards, with media changes following every three days.

#### Drug treatment of organoids and qRT-PCR

Organoids were treated with Fentanyl (74 nM) every other day for 8 days, undergoing 4x fentanyl treatments in total. CEROs were collected in TRIzol (Invitrogen) at various time points throughout maturation. CEROs were subjected to dissociation and lysis in the TRIzol followed by chloroform treatment. The RNA was precipitated from the aqueous phase using isopropanol. Finally, the RNA pellet was washed using 75% ethanol prior to quantification using Nanodrop. Following RNA isolation, cDNA was synthesized by reverse transcription using iScript Reverse Transcription Supermix (Bio-Rad), as per manufacturer’s guidelines. Quantitative real time PCR was performed using SYBR green (Thermo Fisher Scientific, #4309155).

#### Cerebellum organoid cryosection and immunofluorescence

Briefly, the organoids were collected in PBS at multiple timepoints throughout maturation. The organoids were given two PBS washes after which they were subjected to fixation using 4% paraformaldehyde for 20 min at 37°C. The organoids were washed again twice before saturating in 30% sucrose solution overnight at 4°C. CEROs were embedded in Tissue-Tek O.C.T. (Sakura, #62550) blocks and cryo-sectioned into 16 µm thick tissue sections on positively charged glass slides. For immunostaining, the samples were subjected to antigen retrieval using 1X citrate buffer (Sigma) at 90°C in a hot air oven. The slides were then washed thrice for 5 mins each followed by addition of PBS containing 0.1% triton X-100 for 20 mins. Following three washes, post permeabilization the slides were then blocked using 1% BSA in PBST for 30 mins. The samples were treated with primary antibody solution diluted in a blocking buffer at 4°C overnight. The next day, the slides were given three PBS washes and a secondary antibody was added at a 1:1000 dilution for an hour at room temperature. Finally, nuclear stain is added to the sample (DAPI, 1 µg/mL) for 10 mins. The slides were washed in PBS and sealed using an antifade mounting reagent (ProLong diamond Invitrogen).

#### Live calcium imaging of organoids

Fluorescence live cell imaging on the organoids was performed on Krieger microscope with a multiphoton laser equipped to maintain CO_2_ and temperature conditions. For calcium imaging, the organoids were treated with Fluo-4 Direct Calcium Assay Kit (Invitrogen) as per manufacturer guidelines. Briefly, the organoids were washed twice with PBS before addition of Fluo-4, AM, 1000X diluted to 1X in PowerLoad™ concentrate supplied in the kit. The dye was mixed by vortexing and added to the organoids followed by an incubation for 15 mins at 37°C and 30 min at room temperature. Fluo-4, AM loading solution was removed and the organoids were washed once in PBS. Time-lapse images were captured at 494 nm excitation with frames captured at a frame rate of 7 Hz to capture the changes in calcium dependent fluorescence. Images were spatially reduced by 25% using ImageJ (1.54r)^115^ and data analysis was performed using the CaImAn software package (v1.13.1)^116^.

Briefly, the brightest signal across time (max projection) highlighted persistent structures in the overall field of view. Local correlation image (CaImAn local_correlations) represents spatially coherent activity, making neuron-like structures easier to visualize. Further, sample drift and non-uniform motion were corrected so that neural signals remain aligned over time using CaImAn Motion Correct (NoRMCorre-based, piecewise rigid). Neurons were identified and their activity was separated from the background using CaImAn CNMF (constrained nonnegative matrix factorization) which was run in patches to handle local structure and overlap. Key CNMF parameters (e.g., patch size, overlap, expected neuron size, and components per patch) were tuned based on visual inspection of the quilt plot (view_quilt) to ensure patches span multiple neuron diameters while maintaining enough overlap. Signal-to-noise ratio (SNR), spatial correlation (rval), and CNN (convolutional neural network)-based shape classifier were used to retain components that look and behave like neurons and reject noise/artifacts. Components were visually inspected using contour overlays (plot_contours_nb) to verify spatial plausibility of accepted ROIs (default thresholds: min_SNR = 2.0; rval_thr = 0.85; min_cnn_thr = 0.99; cnn_lowest = 0.5). To isolate true calcium transients from technical noise, we employed a percentile baseline. This approach calculates the relative percentage change in fluorescence (DF/F) against a locally determined resting state. We used CaImAn estimates.detrend_df_f with a running percentile baseline to obtain normalized DF/F traces, where DF/F = (F(t)-F_0_)/ F_0,_ and F_0_ is the moving baseline.

To quantify the total calcium activity of individual neurons, we calculated the integrated calcium activity. This was defined as the Area Under the Curve (AUC) of the Delta F/F signal over the entire recording duration, computed using the trapezoidal rule (numpy.trapz). To assess functional output, we calculated the mean firing rate (Hz) for each neuron. Spike probabilities derived from deconvolution were summed and normalized by the total recording duration (35.71 seconds).

To account for the nested structure of the data (individual neurons sampled from distinct organoids), statistical significance was determined using linear mixed-effects models (LMMs). For both integrated calcium activity (AUC) and mean firing rate (Hz), the experimental condition (fentanyl vs. control) was treated as a fixed effect, while the organoid sample ID was treated as a random effect. This approach controls for intra-sample correlation and prevents the inflation of type I error associated with pseudoreplication. All analyses were performed in Python (v3.10+) using the pandas, scipy.stats, and statsmodels libraries. Data visualization was performed using seaborn, with results presented as individual neuronal data points overlaid on boxplots representing the median and interquartile range (IQR).

### 10x chromium single cell RNA sequencing for CERO

The organoids were washed using PBS and collected in a 24 well ultra-low attachment plate; one organoid per well. CEROs were treated with Accumax and kept at 37°C, at 150 rpm for 15 mins followed by pipetting using a 1 mL tip. The organoids were put back on the shaker (150 rpm) for an additional 10 min at 37°C. The cell clumps are collected in a microfuge tube, and spun at 500 g for 10 min. The cell supernatant was removed, and the pellet was resuspended in a suitable buffer, PBS, or HBSS. Finally, the cells were passed through a 37 μm filter and single cell suspension was obtained. Single cells are counted using a haemocytometer and an automatic cell counter. Dead cell removal was performed for samples that had less than 70 % live cell population.

ScRNA-seq sequencing was performed as per 10X guidelines for Next GEM single cell 3ʹ and GEM-X 3’. A total of 10 organoids samples; 6-control, 4-fentanyl treated CEROs were sequenced. Briefly, approximately 10,000 cells per sample were loaded onto a Chromium Single Cell Chip to be processed into single-cell gel beads in emulsion (GEMs). The sample libraries were pooled based on molar concentrations and sequenced on a NovSeq instrument (Illumina).

Raw RNA sequencing data were processed using Cell Ranger Count (v9.0.1) using Human (GRCh38) 2024-A as reference. Ambient RNA removal was performed using SoupX (v1.6.2) and doublet removal was done using DoubletFinder (v 2.0.4). Downstream analysis of filtered data was done using Seurat (v 5.2.1). Briefly, normalization, scaling, and variance stabilization were performed using SCTransform for individual samples. Samples were integrated using Harmony (v1.2.3). Cells were clustered using the Louvain algorithm and visualized using UMAP. Cells were annotated based on canonical marker gene expression. Differential gene expression analysis was performed using pseudo bulked cell profiles per sample for fentanyl treated vs control samples using DESeq2^105^ in Seurat.

### Spatial Transcriptomics

#### Xenium sample prep and assay

In situ gene expression was assayed using Xenium slides and reagents. Briefly, fresh frozen PFC tissue was embedded in OCT, sectioned in 10 µm slices, and placed on the Xenium slides. Tissue preparation and sectioning was done at the Sanford Burnham Prebys Histology Core. Subsequent sample processing was carried out at the Spatial and Functional Genomics platform, Center for Epigenomics, UC San Diego according to manufacturer instructions. Gene list consisted of 266 genes from the Xenium Human Brain panel, 91 genes from top DEGs identified via snRNA-seq, and 9 HIV genes. The HIV gene list encompassed the viral structural (*gag* and *env*), enzymatic (*pol*), and regulatory (*vif*, *vpr*, *vpu*, *tat*, *rev*, and *nef*) genes. HIV gene probes were designed based on the HXB2 assembly.

#### Data analysis

Xenium data were analyzed using Seurat^100^. Cells with less than 3 transcripts were removed. Data were normalized and variance stabilized using SCTransform. The first 30 PCs were used to reduce data dimensionality. Cells were clustered using the Louvain algorithm and communities were visualized using UMAP. Cell type annotation was carried out via label transfer using RCTD with the PFC snRNA-seq data from this study as the reference. Annotations were then projected onto spatial images to visualize cortical layering and cell type distribution in the slices. Tissue sections were divided into white matter and grey matter regions based on the boundary between cortical neuron layers and oligodendrocytes. Next, the difference in the expression of glial reactivity and inflammation associated genes in glial cells between white and grey matter was assessed. First, white and grey matter regions in HIV+ samples were subsampled next to the separation boundary such that a roughly equivalent number of glial cells was found in each region. Differential gene expression was tested using the FindMarkers function in Seurat with the Mann Whitney U test (FindMarkers param test = “wilcox”).

## Acknowledgements

We are grateful to Drs. John Satterlee, Howard Fox, and Susan Morgello for their valuable insights and suggestions regarding brain tissue selection and for their guidance throughout these studies. We thank Dr. Peter Strick for helpful discussions and advice on cerebellar anatomy and function. We also thank Dr. Kristen Jepsen of the Institute of Genomic Medicine at UC San Diego for assistance with HTSeq. Immunofluorescent images of organoids were acquired with support from the Stem Cell Genomics Core at the Sanford Stem Cell Institute. Live cell imaging of organoids was performed at the Nikon Imaging Center at UC San Diego. We thank Dr. Peng Guo and Dr. Richard Sánchez for their assistance with confocal microscopy experiments. Spatial genomics experiments were carried out at the Center for Epigenomics at UC San Diego, which is supported in part by the UC San Diego School of Medicine. We are grateful to Jesus Flores, and Eric Boone for their technical contributions to data generation using the spatial genomics platform. We also thank Guillermina Garcia at the Histology Core Facility, Sanford Burnham Prebys, for tissue sectioning in support of the spatial genomics experiments. We are grateful to the members of the SCORCH Consortium for their valuable discussions and contributions throughout the course of this work. Sequencing data were generated at the UC San Diego IGM Genomics Center using an Illumina NovaSeq 6000 instrument acquired with support from a National Institutes of Health Shared Instrumentation Grant (S10 OD026929). This work was supported by funding from: (1) NIH grants U01DA053630 and R01DA049524; (2) The John S. Dunn Foundation for the UTHealth Houston Brain Collection; (3) The James B. Pendleton Charitable Trust; (4) The Translational Virology Core at the San Diego Center for AIDS Research (P30 AI036214), and by NIH grant AI169609 (P01 Smith HOME). In addition, this work was made possible by shared resources funded through the NIMH, NIA, NIDA, and NINDS under the following contracts: Texas NeuroAIDS Research Center (TNRC): 75N95023C00016; California NeuroAIDS Tissue Network (CNTN): 75N95023C00014; National Neurological AIDS Bank (NNAB): 75N95023C00017; Manhattan HIV Brain Bank (MHBB): 75N95023C00015; Washington University in St. Louis School of Medicine (WUSM): 75N95024C00027; and Data Coordinating Center (DCC): 75N95023C00013. Additional support for data deposition was provided by NIDA award UM1DA052244. The content of this publication is solely the responsibility of the authors and does not necessarily represent the official views of the NNTC or the National Institutes of Health.

## Author Contributions

TMR, KJG, EAM and AW conceived the study, designed experiments and supervised the study. YL, TV, PK, HM, AG, and XC processed sequencing data. PK generated the combined HIV-Human reference genome. KJG, EAM, AG, TV, XC, EG, and AH designed and conducted the bioinformatics analyses on multiome data. DJM, SG, DMS, and CW-B assisted in donor selection and obtaining brain tissues. SJ, GB, and AG assisted in brain donor selection and SJ dissected the brain samples. SKT, SJ, AW and JB led the protocol development for multiome sequencing from the human brain. JB performed nuclei isolation and generated the snRNA-seq and snATAC-seq libraries. SJ generated the organoid model and performed the fentanyl administration, with SJ GB TV and AG performing analyses. SJ and GB designed spatial transcriptomics experiments and conducted data analysis. QZ assisted in experimental design and data analysis of spatial transcriptomics experiments. EG performed caQTL analyses and AH XC ran SCENIC+. XC developed the genome browser and UCSC cell browser and the landing page for all interactive tools. TMR, EAM, KJG, AW acquired funding and initiated the study. TMR, KJG, EAM, AW, AG, TV, XC, SJ, GB, and EG contributed to writing the manuscript. All authors approved of and contributed to the final version of the manuscript.

## Data Availability

Data was produced as part of the Single Cell Opioid Response in the Context of HIV consortium (SCORCH: RRID:SCR_022600). Publicly accessible data is available at NeMO Archive (RRID:SCR_002001) under identifier nemo:col-c9h2ph2 (https://assets.nemoarchive.org/col-c9h2ph2). Access to all protected raw data associated with this study is managed by dbGaP and can be requested with the identifier phs004026.v1.p1. Processed data are deposited at NeMO Analytics. Codes for the data processing and analysis will be made available at publication. A centralized landing page with links to all interactive resources is available at https://brainome.ucsd.edu/SCORCH.

## Description of Additional Supplementary Files Document

Table S1 Overview of Donor Metadata (e.g. age, HIV status, post-mortem interval)

Table S2 Demographic distribution across conditions and brain regions

Table S3 Cell type marker genes for each brain region

Table S4 SCENIC+ GRN specificity scores (RSS).

Table S5 Differentially expressed genes (DEGs) for glial cell types between brain regions (FDR<0.1)

Table S6 Enrichment of genetic risk for neuropsychiatric diseases within cell type-specific cCREs (LDSC)

Table S7 GWAS summary statistics references for LDSC partitioned heritability results

Table S8 Significant caQTL results for neuronal cell types in each region..

Table S9 Significant motifs enriched in caQTLs using motifBreakR.

Table S10 TF motifs enriched in cell type-specific cCREs (Cell type motifs, Entropy + HOMER)

Table S11 Differentially expressed genes (DEGs) for HIV/Control, OUD/Control, and OUD-HIV interaction analyses (FDR < .25)

Table S12 GSEA Pathway Enrichments (OUD Glia only, HIV, OUD-HIV Interaction)

Table S13 Differentially expressed genes (DEGs) for fentanyl-treated vs untreated organoids

Table S14 Differentially expressed genes (DEGs) for reactive glia compared to non-reactive glia (FDR<0.1)

Table S15 fGSEA pathway enrichments for neurons in OUD/control human postmortem analyses and fentanyl/control organoids.

Table S16 SCENIC+ Prefrontal cortex GRNs

Table S17 SCENIC+ Amygdala GRNs

Table S18 SCENIC+ Cerebellum GRNs

Table S19 SCENIC+ fGSEA enrichments

Table S20 SCENIC+ GRN functions

Table S21 Pathway enrichment results for reactive glia (GSEA)

Table S22 Encephalitic HIV+ donor-associated genes GO Results

Table S23 Differences in expression of key genes in white vs grey matter glia in HIV+ samples

Table S24 Differentially accessible chromatin regions for HIV/Control and OUD/Control DESeq2 results (FDR < 0.25).

## Data Overview

We generated single-nucleus multiomic datasets combining chromatin accessibility (snATAC-seq) and gene expression (snRNA-seq) from 93 postmortem human brain samples, derived from 44 donors. Samples were collected from three brain regions—prefrontal cortex (PFC BA10), cerebellum, and amygdala—and spanned four clinical cohorts: control, HIV, OUD, and comorbid OUD and HIV, yielding approximately 4,000–18,000 nuclei per library. The average number of median ATAC fragments per nucleus was 9,507, and transcription start site (TSS) enrichment ranged from 1.52 to 9.37 (median = 5.28), consistent with high-quality chromatin accessibility data. Gene expression profiling detected 153–4,770 genes (median = 1,644) per nucleus, and a median of 75% of RNA reads mapped specifically to nuclei.

## Extended Data Figures

**Extended Data Figure 1.**
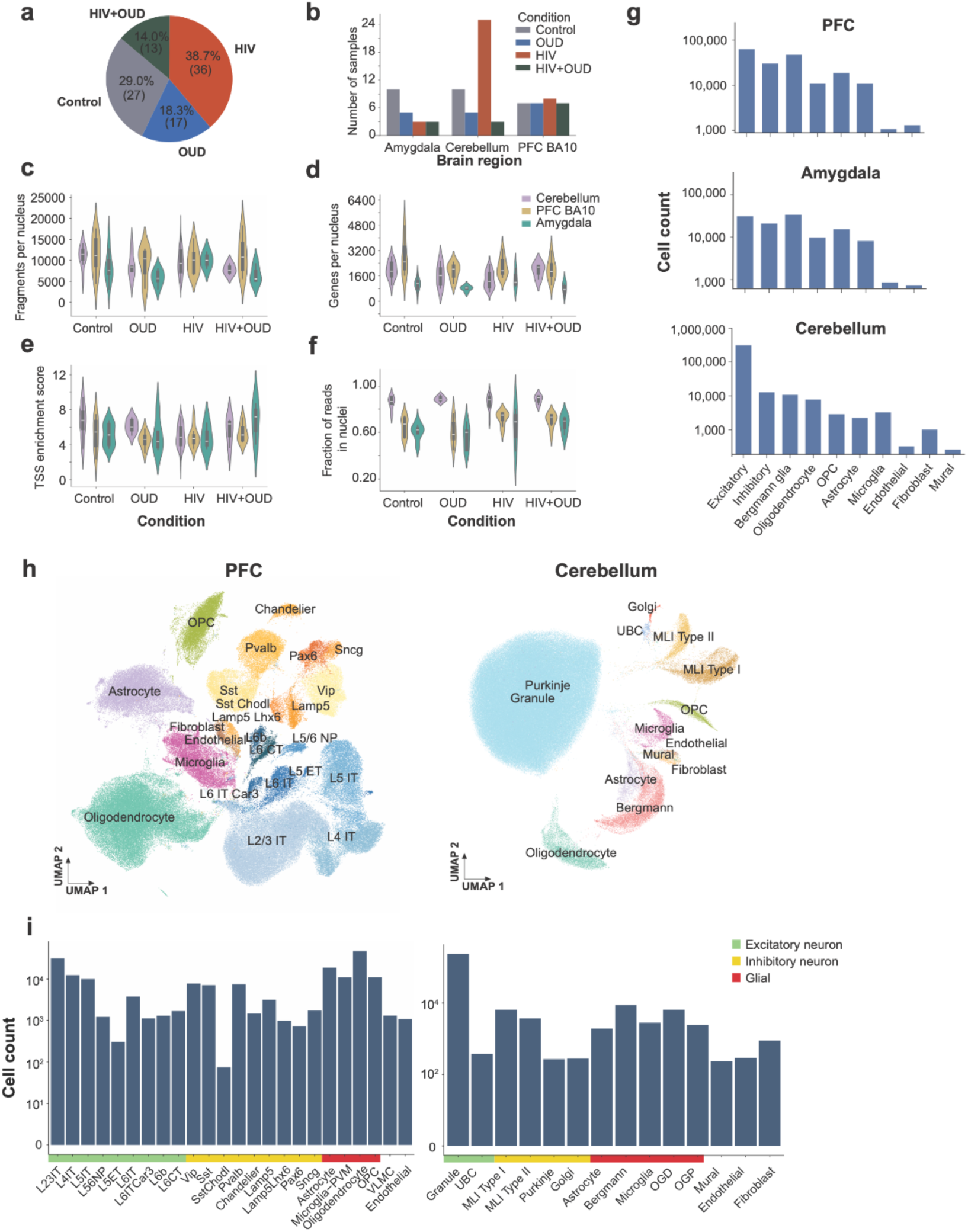
Quality control metrics of single-nucleus multiomic libraries. **(a)** Distribution of samples across clinical conditions: control, HIV-only, opioid use disorder (OUD)-only, and OUD+HIV (n = 93 libraries). **(b**) Number of samples collected from each brain region stratified by clinical condition. **(c-f)**, Quality control metrics for single-nucleus multiomic data grouped by condition and brain region: **(c)** median ATAC fragments per nucleus; **(d)** median number of genes detected per nucleus; **(e)** transcription start site (TSS) enrichment score; **(f)**, fraction of RNA sequencing reads specifically mapping to nuclei. Boxes represent median values and interquartile ranges (IQR); whiskers extend to 1.5× IQR; points denote outliers. Data were derived from 93 postmortem human brain samples obtained from 44 donors. **(g)** Major cell type counts across brain regions. **(h)** UMAPS of PFC and cerebellum labeling neuronal subtypes. **(i)** Number of individual cells assigned to each cell type.

**Extended Data Figure 2.**
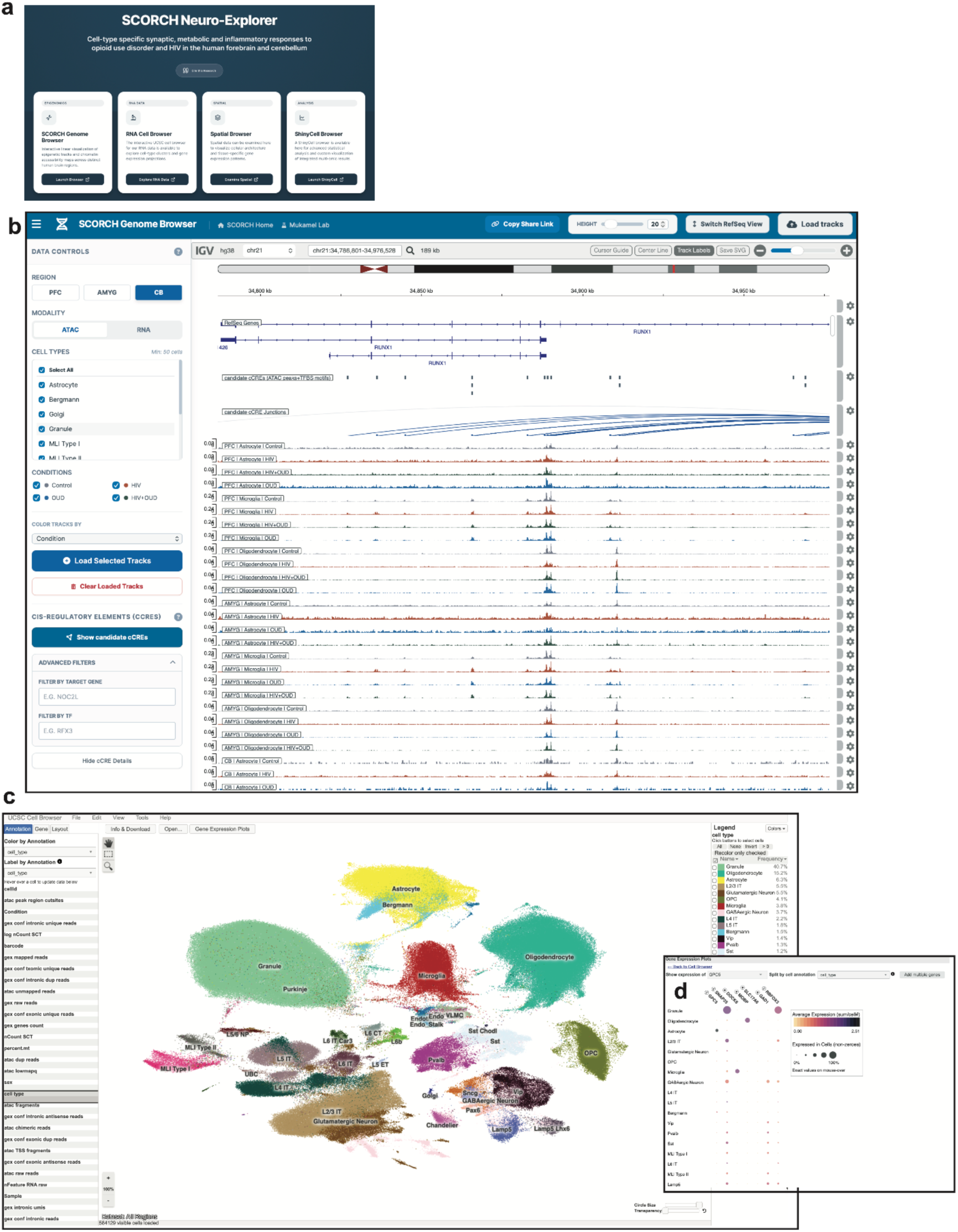
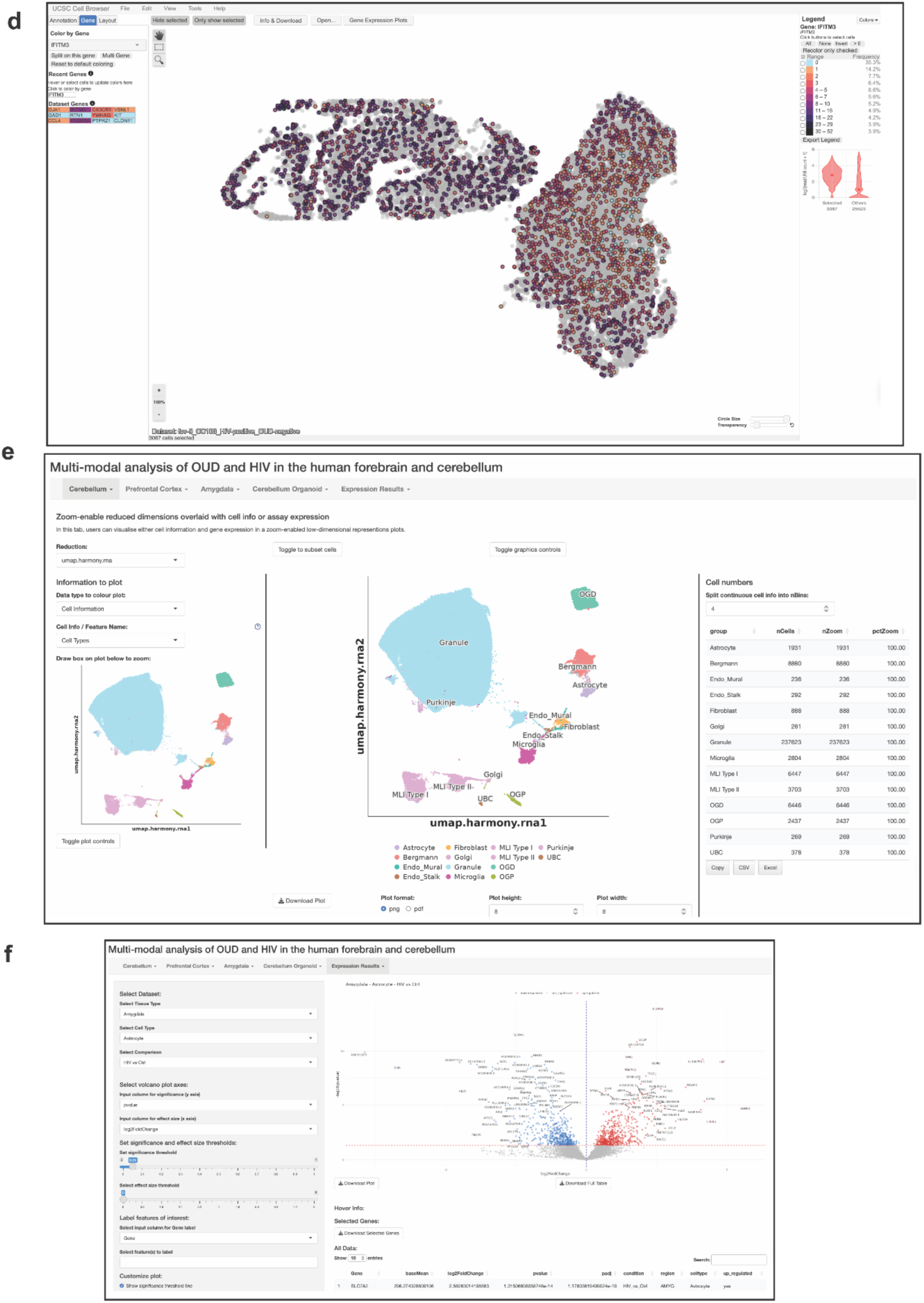
SCORCH Neuro-Explorer web portal for integrated multi-omic visualization across human forebrain, cerebellum and organoid models. **(a)** Landing page of the SCORCH Neuro-Explorer platform providing access to multi-modal datasets from human prefrontal cortex (PFC), amygdala (AMYG), cerebellum (CB) and cerebellar organoids (https://brainome.ucsd.edu/SCORCH). The portal integrates regulatory, transcriptomic and spatial data within a unified analysis environment. (**b)** Genome browser interface for visualization of gene expression, chromatin accessibility, cCREs, gene annotations and condition-specific tracks (https://brainome.ucsd.edu/xic007/SCORCH_Genome_Browser). Users can select region, cell type and diagnostic group to dynamically render comparative accessibility profiles, enabling cross-region and cross-condition inspection of regulatory landscapes. **(c)** UCSC Single Cell Browser for integrated single-nucleus RNA-seq datasets across PFC, AMYG, CB and cerebellar organoids (https://brainome.ucsd.edu/xic007/SCORCH_Cell_Browser/cbBuild). The interface includes (1) dataset selection across regions and organoid models; (2) gene search functionality; (3) UMAP embeddings colored by cell type or metadata; and (4) a dot plot module summarizing gene expression across selected cell types, displaying both mean expression and fraction of expressing cells. **(d)** UCSC Single Cell Browser for spatial transcriptomic datasets (https://brainome.ucsd.edu/xic007/SCORCH_Spatial_Browser/cbBuild). Fields of view (FOVs) are displayed with cells mapped to anatomical coordinates and colored by normalized expression of a selected gene, enabling direct visualization of spatial heterogeneity within intact tissue architecture. **(e)** ShinyCell browser for interactive dataset exploration (http://tools.cmdga.org:3838/scorch_browser). The interface allows metadata filtering (for example, region, condition, donor and cell type), visualization of integrated embeddings and dynamic summaries of cell-type composition across diagnostic groups. **(f)** ShinyCell differential expression module. Volcano plots display gene-level comparisons between selected conditions within defined cell types, with adjustable Log2FC and false discovery rate thresholds. Genes of interest can be highlighted and exported for downstream analysis.

**Extended Data Figure 3.**
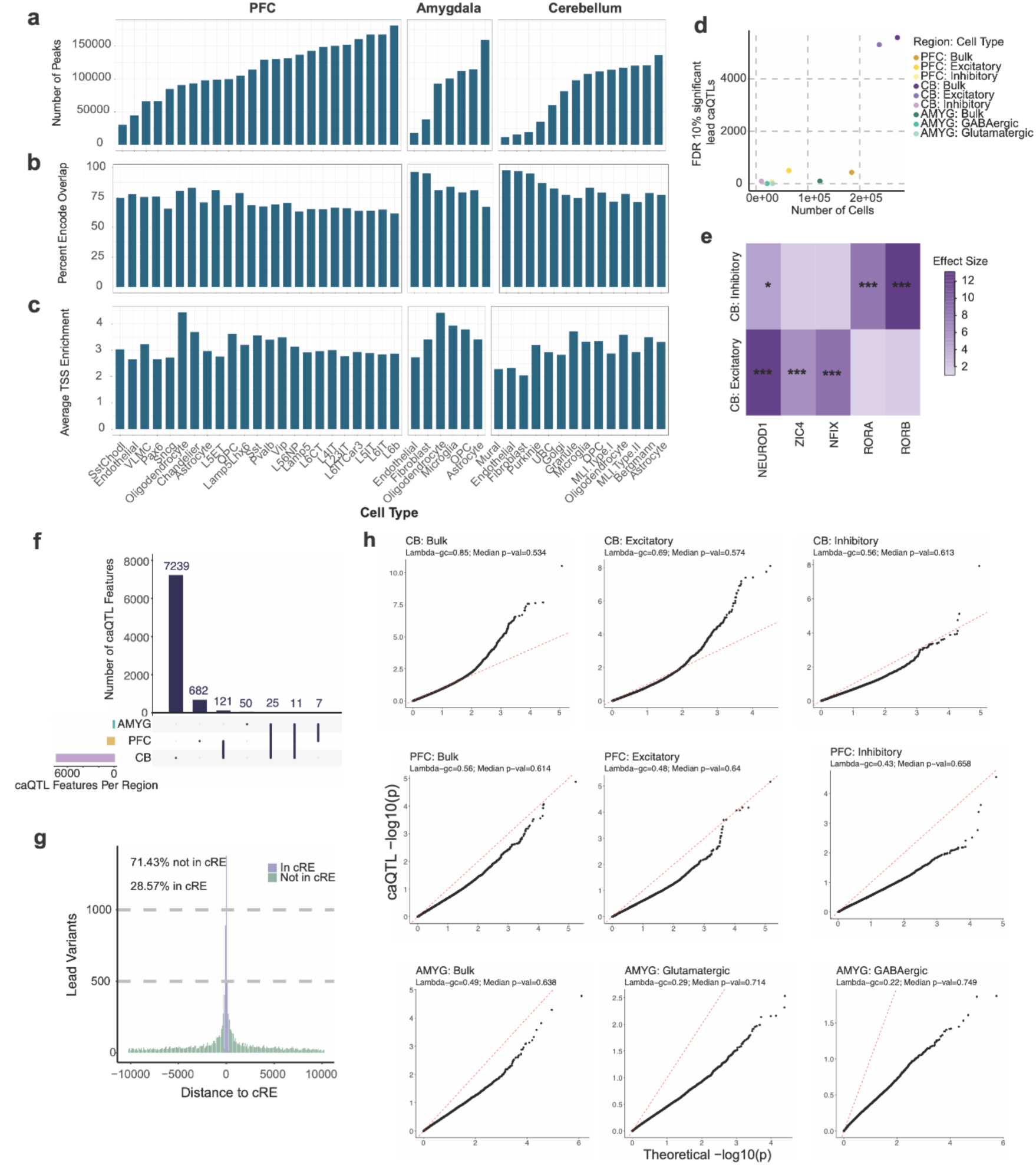
ATAC-seq overview. **(a)** Number of total peaks called per cell type, ordered from lowest to highest number of peaks across cell types in a brain region. **(b)** Percent overlap of called peaks with bulk Encode cis-regulatory elements **(c)** Average transcription start site enrichment (TSSe) for barcodes assigned to the cell type. **(d-h)** Overview of Quantitative Trait Loci (QTLs). (**d**) Number of significant lead (most significant in the locus) caQTL variants as a function of the number of cells for each cell type analyzed. (**e**) Shared and cell type-specific motifs enriched in caQTLs, *=FDR < 0.10, ** = FDR < 0.01, *** = FDR< 0.001. (**f**) Shared caQTL features across brain regions. (**g**) Distribution of the distance of caQTL variants to cCREs. h): q-q plots for all tested caQTL variants across all cell types analyzed. Dashed, red line at x=y.

**Extended Data Figure 4.**
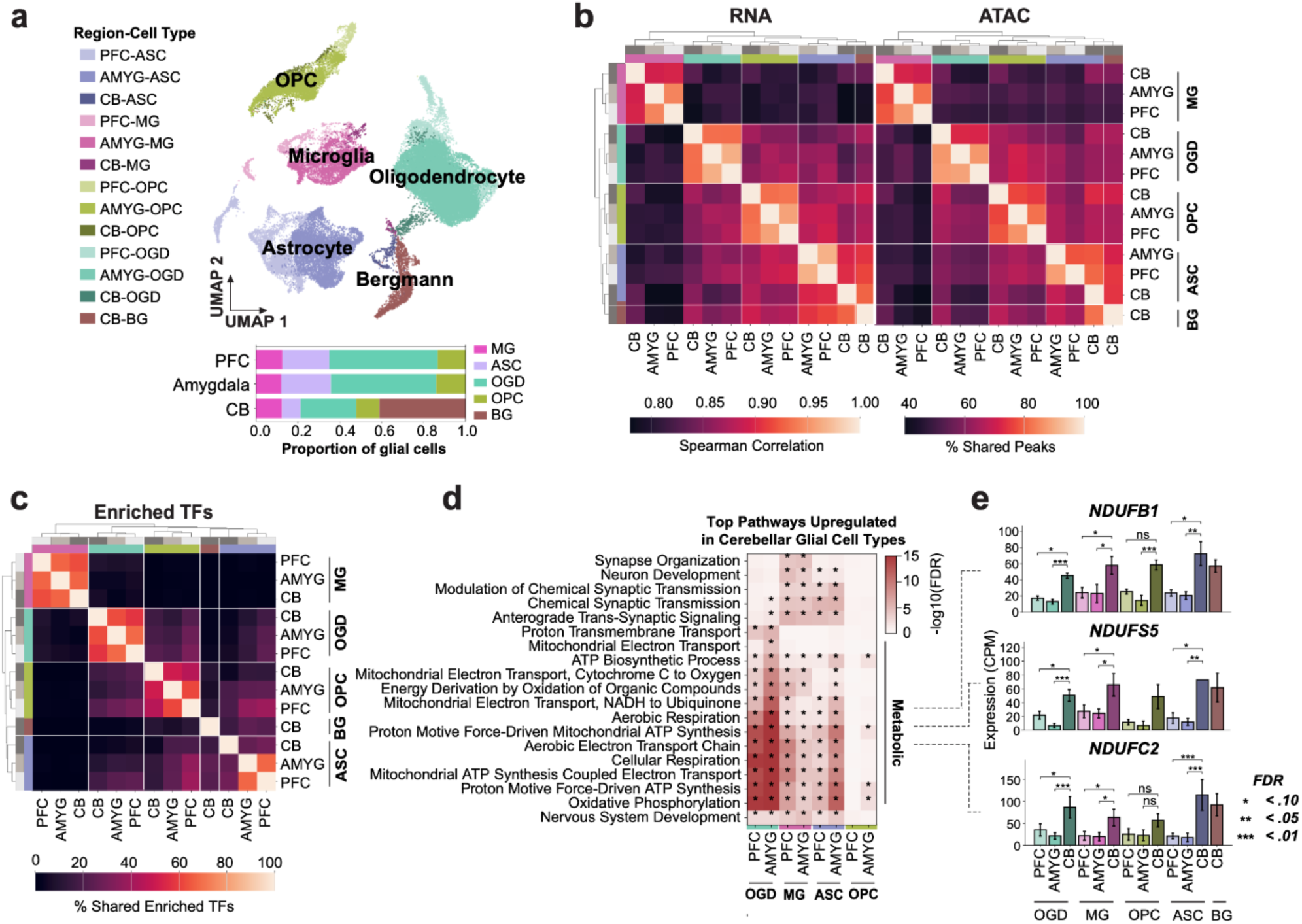
Upregulation of energy metabolism pathways in cerebellar glial cell types compared to PFC and amygdala. **(a)** RNA-based UMAP of glial cell types across brain regions and glial cell type counts across regions in control donors (top). Glial cell type proportions across regions in all donors (bottom). ASC, Astrocyte; MG, Microglia, OPC, Oligodendrocyte precursor cell; OGD, Oligodendrocyte; BG, Bergmann glia. (**b**) Spearman correlation heatmap between glial cell types and brain regions from RNA data in control donors (left) and percentage of shared peaks between glial cell types and brain regions from ATAC data in all donors (right). (**c**) Percentage of shared significant enriched transcription factors (TFs) between cell types and regions. (**d**) Top 50 GO enriched pathways in the cerebellum, derived from differential gene expression analyses between brain regions in glial cell types. (**e**) Expression of the top 3 genes most highly shared between 90 significantly upregulated metabolic pathways in the cerebellum compared to the PFC and amygdala (donor pseudobulk from controls with tissue from all 3 regions). Error bars show standard error.

**Extended Data Figure 5.**
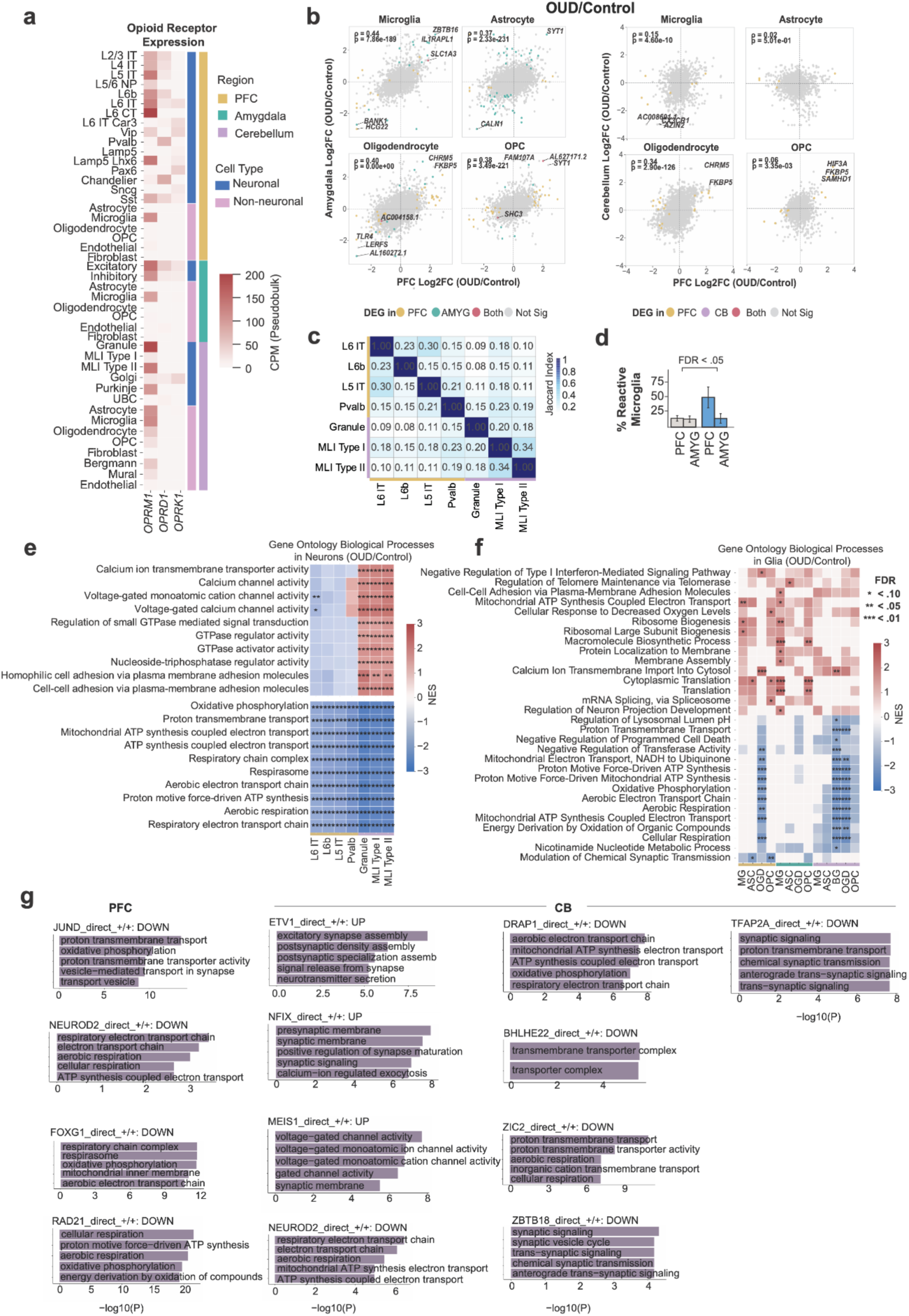
(a) CPM normalized expression of opioid receptors across all cell types in the PFC, amygdala, and cerebellum. **(b)** Log2FC (OUD/Control) correlation results between the PFC and amygdala (left) and the PFC and cerebellum (right) across glial cell types. **(c)** Jaccard index values for DEGs shared across seven transcriptionally impacted neurons. **(d)** Reactive microglia proportion across and conditions in the PFC and amygdala (Linear Mixed Effects Model, FDR correction). Error bars show standard error. **(e)** Full pathway names for gene ontology pathway results shown in Fig. 2c **(f)** Gene ontology results in glia across brain regions (OUD/Control). **(g)** Target gene enrichments for GRNs significantly altered in OUD (FDR<0.1) not shown in Fig. 2d, if there are significant pathway enrichments for that GRN target gene set (FDR < 0.05).

**Extended Data Figure 6.**
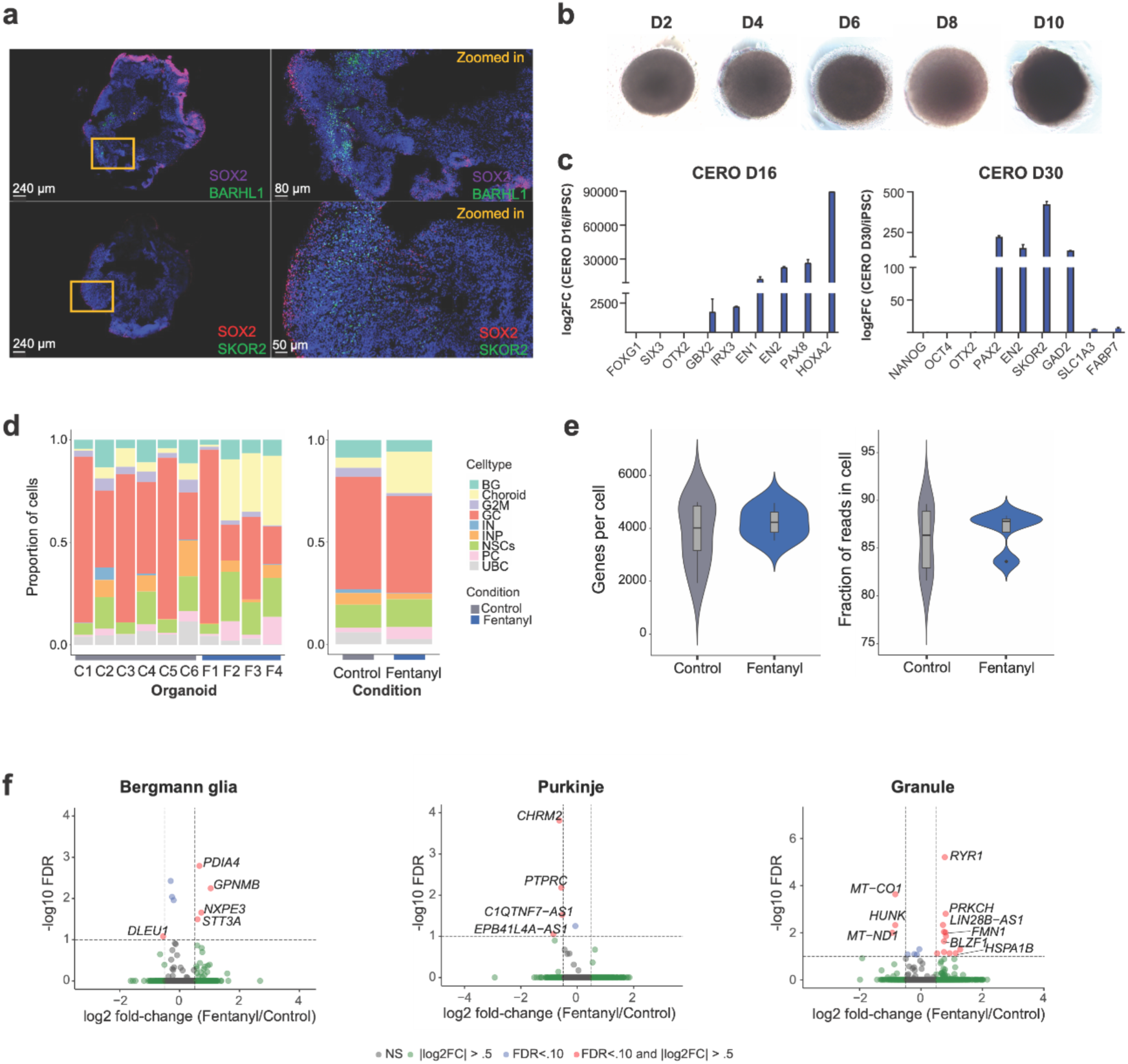
CERO development and scRNA-seq. (**a**) Immunofluorescence staining of BARHL2+ GC progenitors and SKOR2+ inhibitory newborn Purkinje cells in 1 month old CERO. SOX2 and Nestin identify neural progenitors. The nuclei were stained with DAPI (blue). (**b**) DIC images illustrate typical morphology of cerebellar organoids on different days of CERO development. **(c)** qRT-PCR of region-specific markers on day 16 (left) and day 30 (right) CERO. Organoid patterning unique to cerebellum development was observed. Error bars show standard deviation. (**d)** Bar plots showing the proportion of cell types by organoid (left) and condition (right). (**e)** CERO sc-RNAseq QC metrics. Genes and fraction of reads mapped to cell-associated barcodes, a measure of ambient RNA, across conditions. Inner box plots show median and IQR. (**f)** Volcano plots showing differentially expressed genes in Bergmann glia, Purkinje cells, and granule cells after fentanyl treatment. |log2FC|>0.5, FDR < 0.1 (grey: not significant; blue: significant FDR; green: significant log2FC; red: significant FDR and significant log2FC).

**Extended Data Figure 7.**
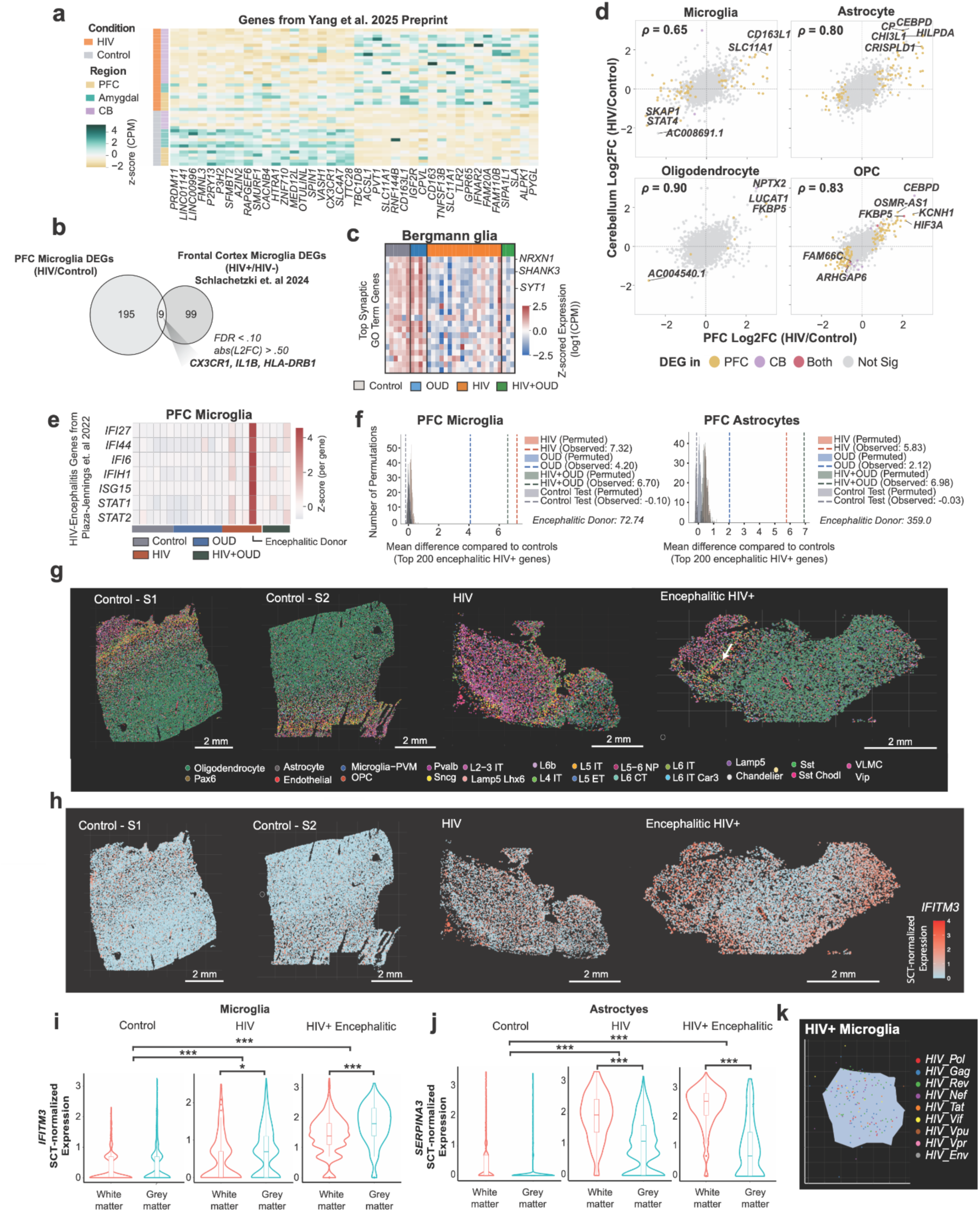
(a) Heatmap of top up and downregulated DEGs in microglia from Yang et. al 2025 Preprint plotted in our PFC microglia data. **(b)** DEGs that overlap in our PFC microglia data compared to Schlachetzki et. al 2024’s HIV+ microglia vs HIV- microglia. **(c)** Most common genes in Synaptic GO pathways from Fig. 2b plotted in Bergmann glia. Each column shows a pseudobulk expression profile for Bergmann glia from one sample. (**d**) Log2FC Spearman correlation results between the PFC and cerebellum across glial cell types. All correlations p-val < .001. **(e)** Expression of HIV-encephalitis related genes in PFC microglia (donor pseudobulk). **(f)** Genes that are strongly upregulated in the HIV+ encephalitic donor are also upregulated in non-encephalitic disease conditions. X-axis shows the mean Log2FC for each group relative to control donors, averaged across the top 200 genes with greatest median difference in expression in encephalitic HIV+ donor compared to controls in microglia (left) and astrocytes (right). Histograms show distribution of Log2FC differences for randomly permuted gene sets. **(g)** Spatial transcriptomics (Xenium) showing annotated cell types in control, HIV, and encephalitic HIV+ donors. **(h**) Spatial transcriptomics *IFITM3* expression. (**i**) Microglia *IFITM3* expression in white vs gray matter in each image. *=FDR < 0.10, ** = FDR < 0.01, *** = FDR< 0.001. (**j**) Astrocyte *SERPINA3* expression in white vs gray matter in each image. **(k)** Spatial configuration of transcripts within a single HIV+ microglial cell from the HIV+ encephalitic donor (white arrow in panel g).

**Extended Data Figure 8.**
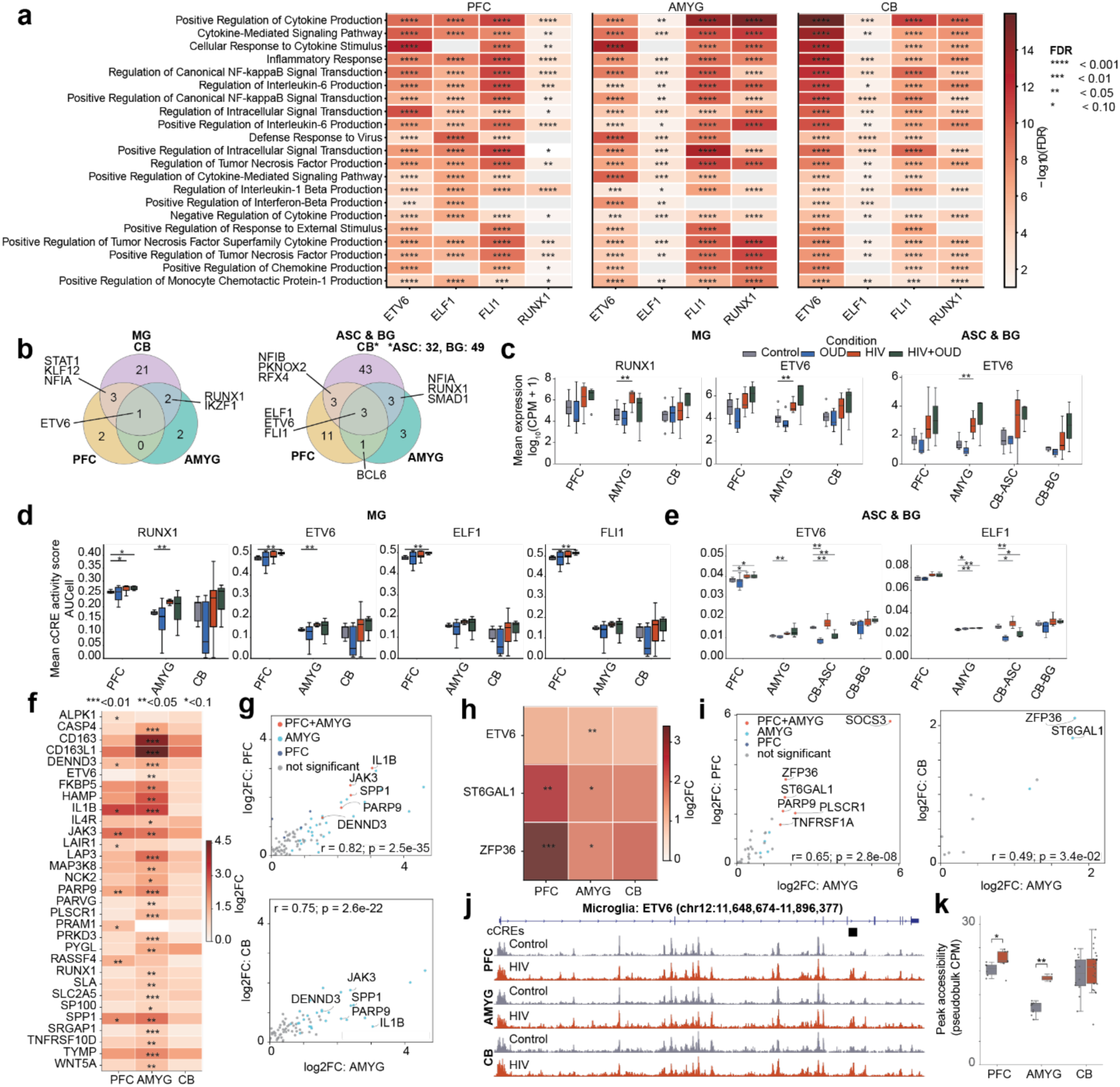
Brain region- and cell-type-specific features of HIV-associated GRNs. **(a)** Gene ontology enrichment of HIV-associated GRN targets. Heatmap shows the top 20 enriched biological process terms (FDR < 0.10) for selected GRNs (ETV6, ELF1, FLI1, and RUNX1) across PFC, amygdala, and cerebellum. Color intensity represents –log₁₀(FDR); grey indicates absence of enrichment. **(b)** Venn diagrams showing cross-regional overlap of HIV-associated GRNs. In microglia, ETV6 is conserved (left). In astrocytes and Bergmann glia, shared enrichment was observed for ETV6, ELF1, and FLI1, indicating parallel but partially distinct neuroimmune regulatory circuits across regions (right). **(c)** Expression of HIV-associated transcription factors in microglia, astrocytes, and Bergmann glia. Boxplots show sample-level log₁p(CPM) expression of ETV6 across microglia, astrocytes, and Bergmann glia, and RUNX1 specifically in microglia, profiled in PFC, amygdala, and cerebellum across Control, OUD, HIV, and OUD+HIV conditions. Boxes represent the interquartile range (IQR), whiskers represent 1.5 IQR, and the horizontal line indicates the median. **(d-e)** HIV-associated GRN activity in microglia, astrocytes, and Bergmann glia. (**d**) Region-based GRN activity (AUCell) for RUNX1, ETV6, EF1, and FLI1 GRNs across PFC, amygdala, and cerebellum in microglia. Boxplots show mean single-cell region activity scores stratified by condition (Control, OUD, HIV, OUD+HIV). (**e**) Region-based GRN activity (AUCell) for ETV6 and ELF1 GRNs across PFC, amygdala, and cerebellum in astrocytes and Bergmann Glia. **(f)** Target region regulation of HIV-associated GRNs in microglia. Heatmap depicts Log2FC (HIV/control) of significant RUNX1 and ETV6 target genes. **(g)** Target gene Log2FC (HIV/Control) between brain regions, highlighting shared significantly regulated targets. **(h)** Target gene regulation of HIV-associated GRNs in astrocytes. Heatmap shows Log2FC (HIV/control) of target genes of ETV6. **(i)** Log2FC between PFC/CB and amygdala for DEG targets. **(j)** Regulatory target gene browser tracks. Genome browser representations show normalized chromatin accessibility (ATAC-seq) at the ETV6 locus in microglia, illustrating a differentially accessible cCRE. **(k)** Boxplot of pseudobulked region accessibility of ETV6 target region stratified by condition (Control, HIV).

**Extended Data Figure 9.**
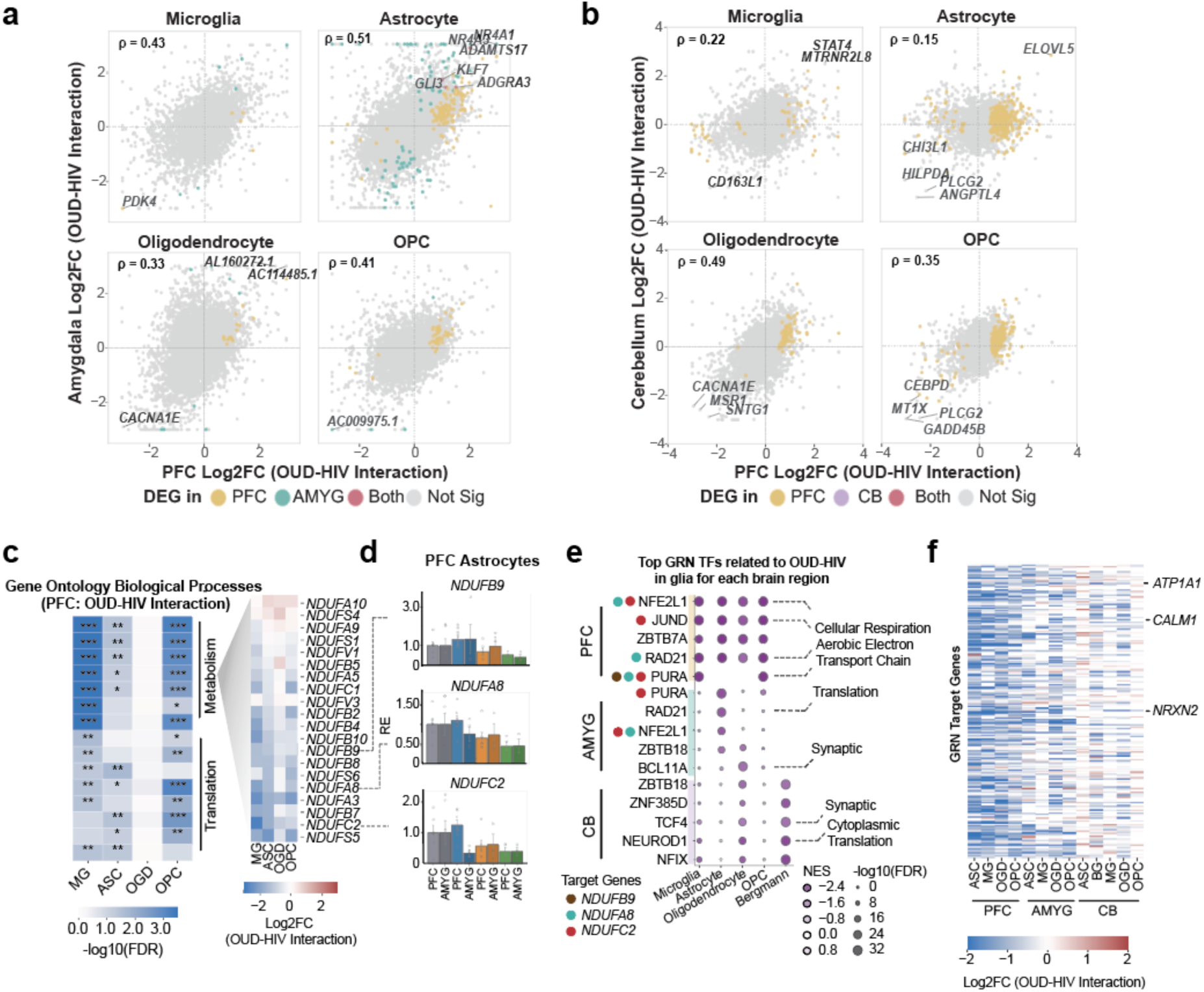
OUD-HIV Interaction impacts on glial cell types. **(a)** Log2FC (OUD-HIV interaction) correlation results between the PFC and amygdala across glial cell types. Genes shown are filtered to those FDR< .95, Log2FC > .10. **(b)** Log2FC (OUD-HIV interaction) Spearman correlation results between the PFC and cerebellum DEGs across glial cell types. **(c)** Gene ontology pathway terms from OUD-HIV interaction differential gene expression analyses in PFC glial cell types (left). Log2FC (OUD-HIV Interaction) of most common genes in metabolic pathways across PFC glial cell types (right). **(d)** Expression of *NDUFB9*, *NDUFA8*, and *NDUFC2*, in PFC and amygdala astrocytes (donor pseudobulk). **(e)** OUD-HIV interaction DEG-enriched GRNs (SCENIC+). **(f)** Log2FC of target genes in top GRNs across glial cell types and brain regions.

## Supplementary Figures

**Figure S1.**
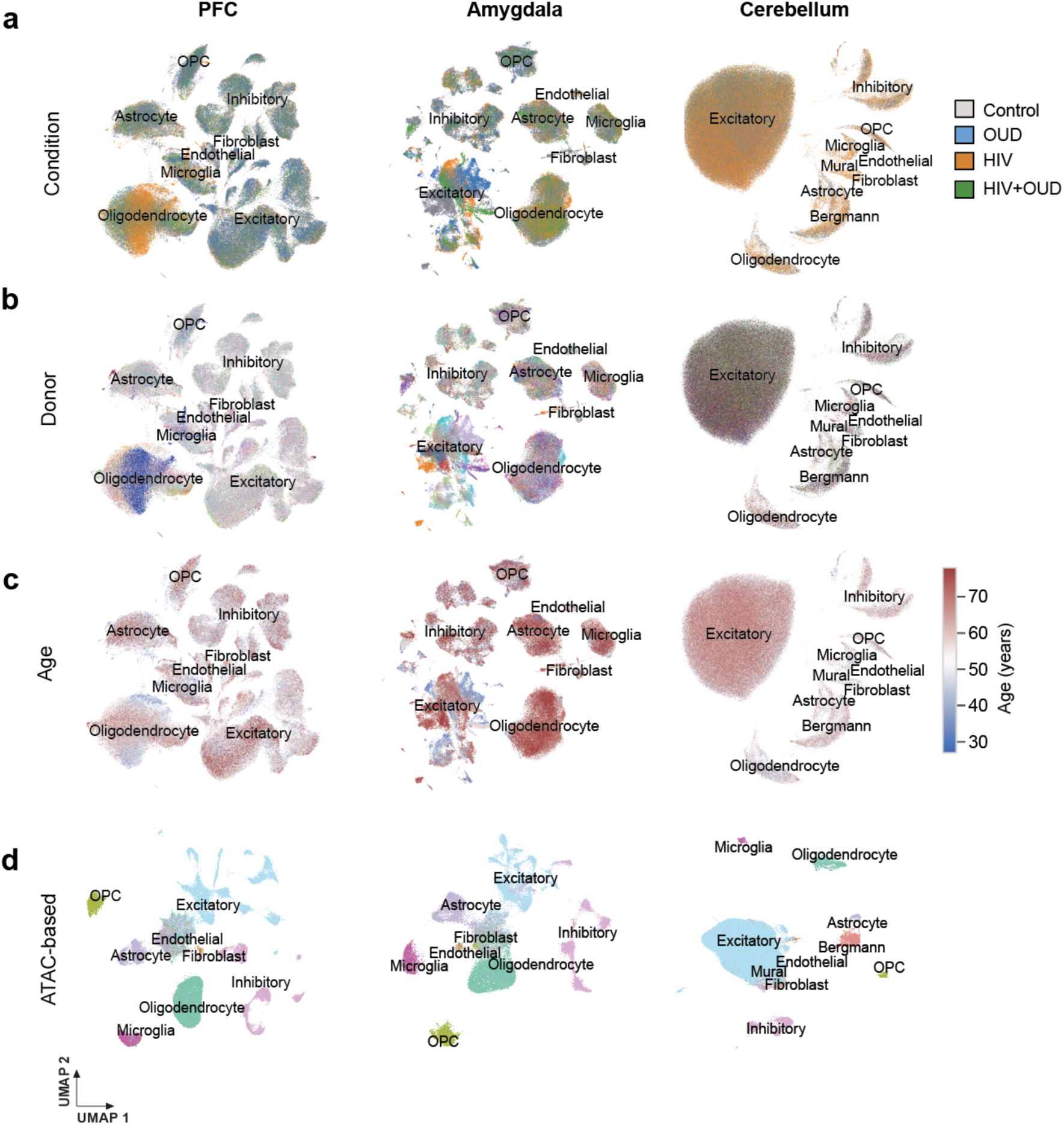
(a-c) UMAPs colored by: (**a**) condition, (**b**) donor, and (**c**) age. (**d**). UMAP clustering based on regional consensus peak set(s).

**Figure S2.**
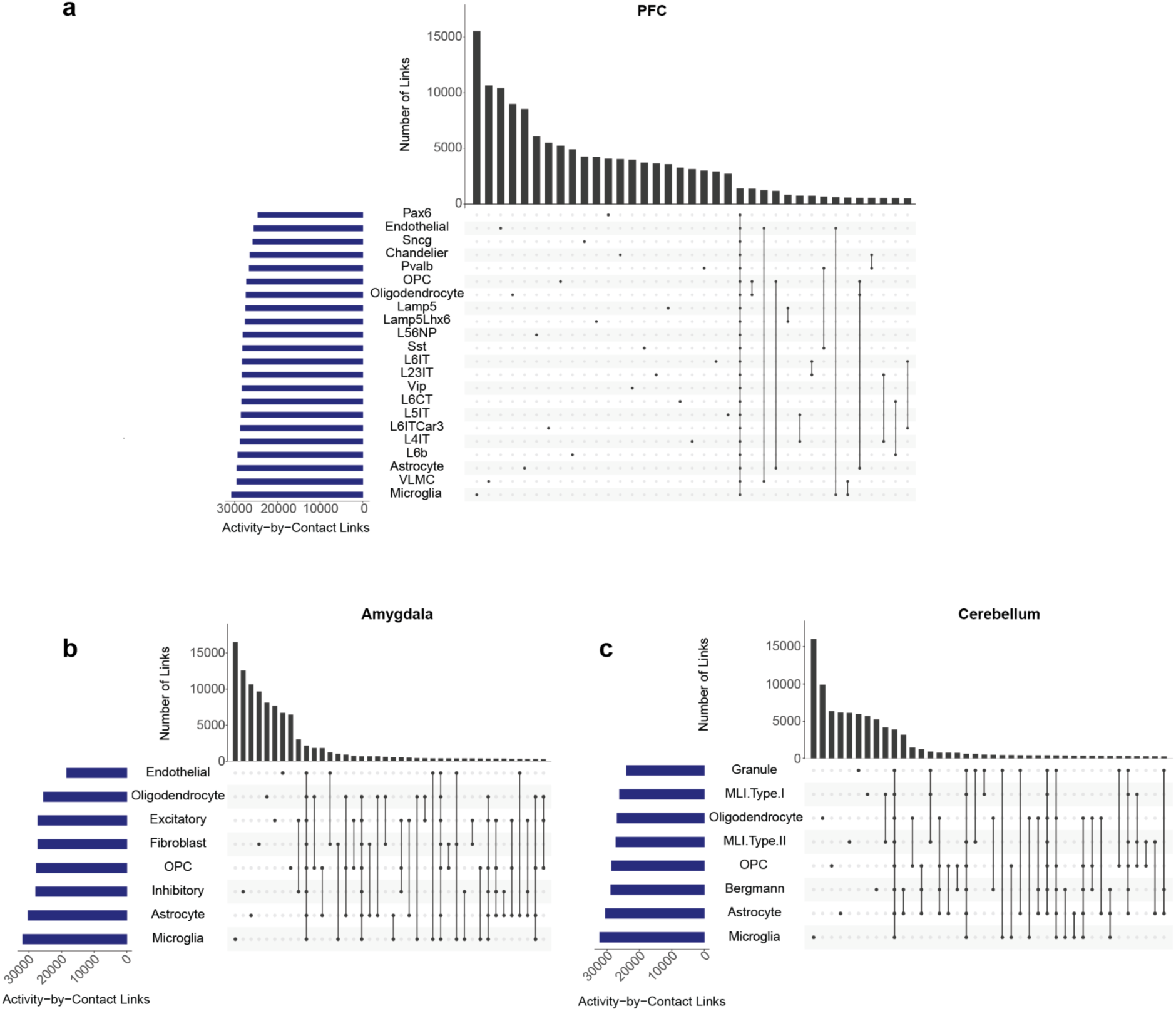
Overview of Activity-by-Contact (ABC) links. Peak-gene links shown for cell types in the **(a)** PFC, **(b)** amygdala, and **(c)** cerebellum.

**Figure S3.**
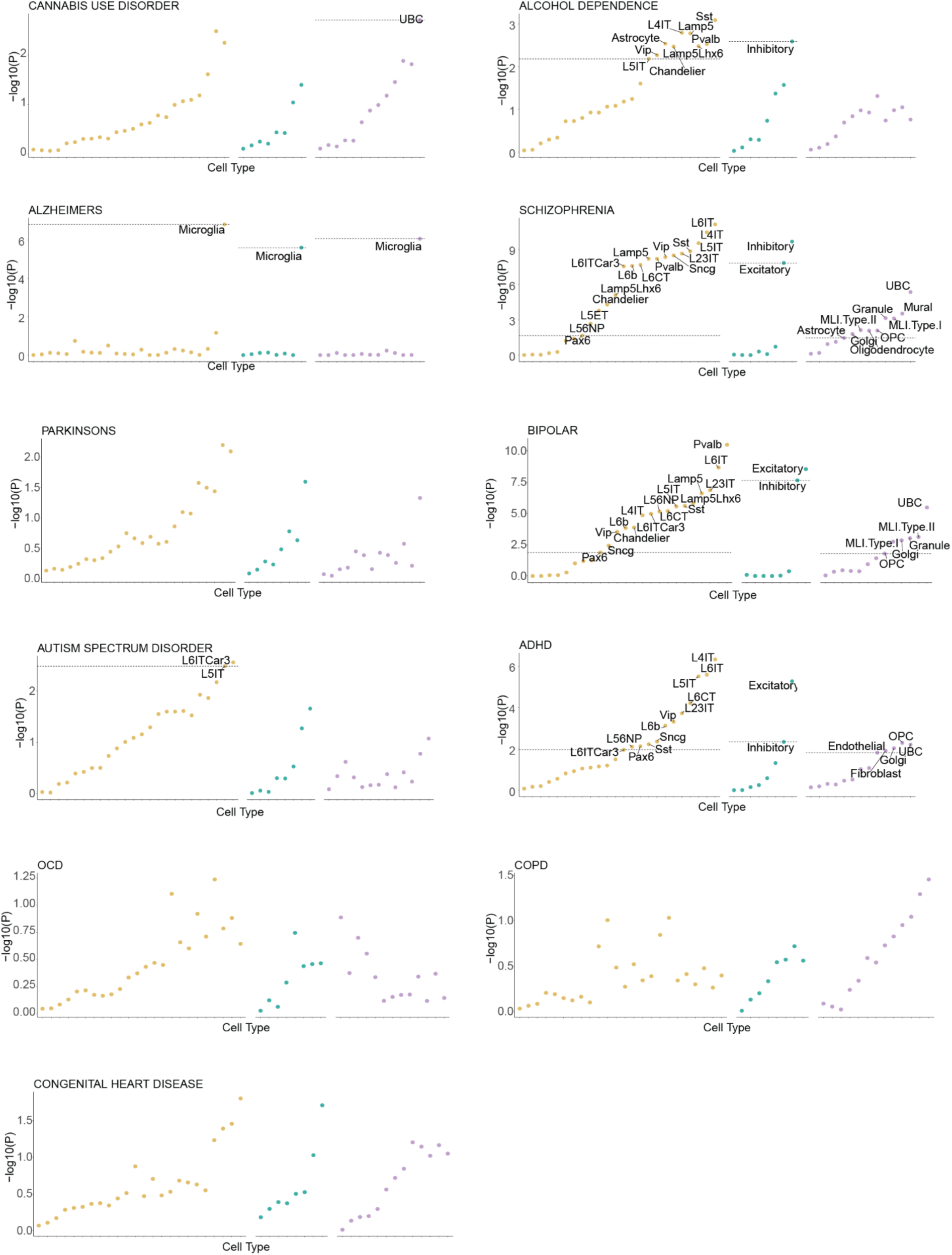
LDSC Cell type significance by trait. Significance of every cell type across regions, for each trait. Raw p-values are shown. Dashed line indicates p-value associated with highest FDR less than or equal to 0.05. Multiple test corrections are performed within each region.

**Figure S4.**
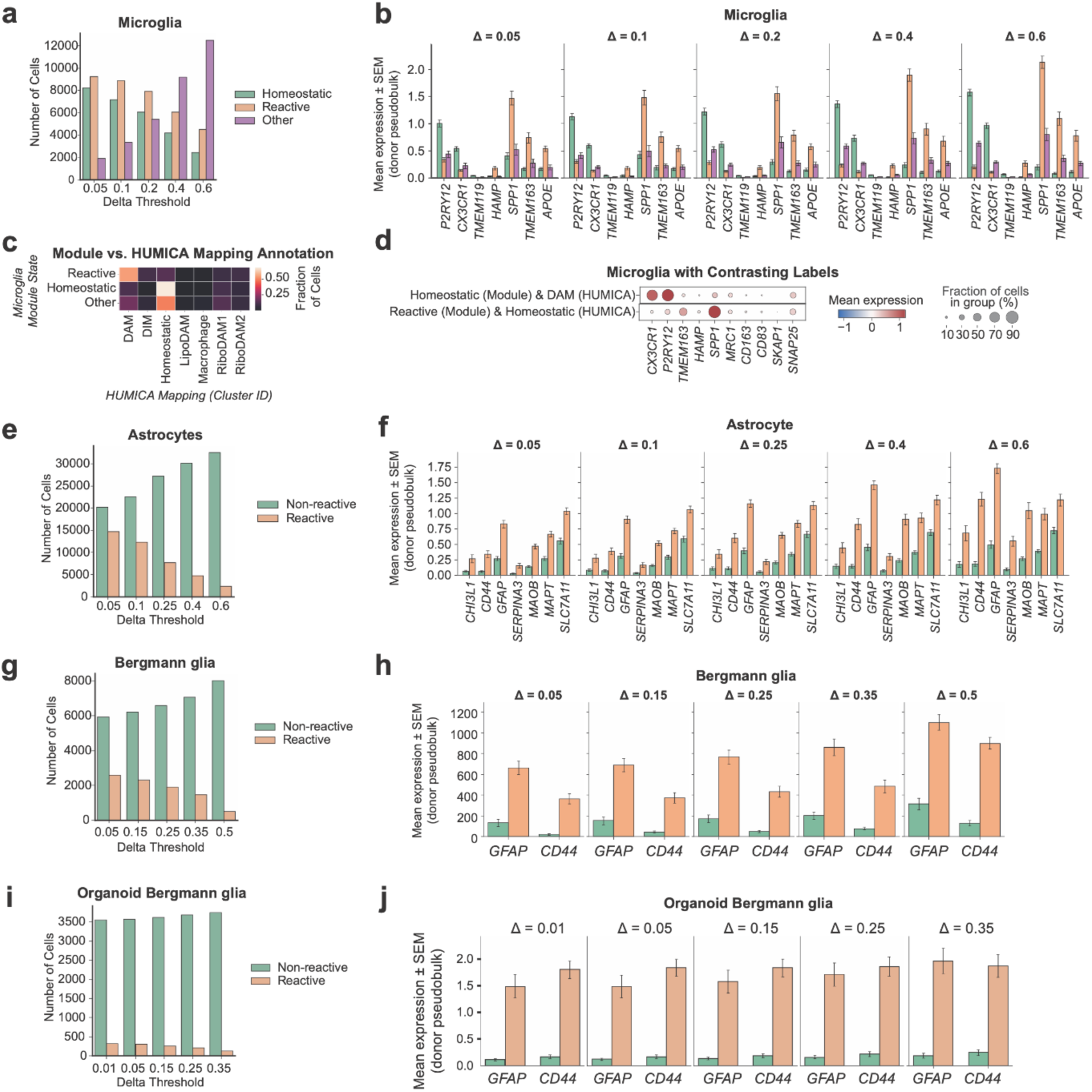
(a) Number of microglia separated into each category according to different delta thresholds. **(b) E**xpression of microglia state marker genes across delta thresholds. Error bars show standard error. **(c)** Confusion matrix showing annotation using mapping to Human Microglia Atlas (HUMICA) or module annotation with marker genes. **(d)** Microglia with opposite annotations according to each method. **(e)** Astrocyte state numbers according to different delta thresholds. **(f)** Expression of astrocyte state marker genes across delta thresholds. **(g)** Bergmann glia state numbers according to different delta thresholds. **(h)** Expression of Bergmann glia state marker genes across delta thresholds. **(i)** Organoid Bergmann glia state numbers according to different delta thresholds. **(j)** Expression of organoid Bergmann glia state marker genes across delta thresholds.

